# Hippocampal theta bursting and waveform shape reflect CA1 spiking patterns

**DOI:** 10.1101/452987

**Authors:** Scott Cole, Bradley Voytek

## Abstract

Brain rhythms are nearly always analyzed in the spectral domain in terms of their power, phase, and frequency. While this conventional approach has uncovered spike-field coupling, as well as correlations to normal behaviors and pathological states, emerging work has highlighted the physiological and behavioral importance of multiple novel oscillation features. Oscillatory bursts, for example, uniquely index a variety of cognitive states, and the nonsinusoidal shape of oscillations relate to physiological changes, including Parkinson’s disease. Open questions remain regarding how bursts and nonsinusoidal features relate to circuit-level processes, and how they interrelate. By analyzing unit and local field recordings in the rodent hippocampus, we uncover a number of significant relationships between oscillatory bursts, nonsinusoidal waveforms, and local inhibitory and excitatory spiking patterns. Bursts of theta oscillations are surprisingly related to a decrease in pyramidal neuron synchrony, and have no detectable effect on firing sequences, despite significant increases in neuronal firing rates during periods of theta bursting. Theta burst duration is predicted by the asymmetries of its first cycle, and cycle asymmetries relate to firing rate, synchrony, and sequences of pyramidal neurons and interneurons. These results provide compelling physiological evidence that time-domain features, of both nonsinusoidal hippocampal theta waveform and the theta bursting state, reflects local circuit properties. These results point to the possibility of inferring circuit states from local field potential features in the hippocampus and perhaps other brain regions with other rhythms.

## Introduction

Oscillations are one of the most prominent features of neural field potential recordings (Buzsáki and Draguhn, 2004; Cohen, 2017). Consequently, they have been extensively studied for decades and their features are known to relate to physiological processes, pathologies, and behavior (Buzsáki and Wang, 2012; Cohen, 2017; Klimesch, 1999; Uhlhaas and Singer, 2010; Ward, 2003; Womelsdorf et al., 2014). Throughout this research, clear evidence has emerged showing that local spiking probability is coupled to phases of the oscillatory local field (Li et al., 1952), resulting in theories about how these oscillations may function to aid communication among brain networks (Fries, 2005; Peterson and Voytek, 2018; Roux and Uhlhaas, 2014; Voytek and Knight, 2015). Though we have a good understanding of how current sources summate and manifest as field potential fluctuations (forward model), interpretations of these oscillations is challenging because many different biological processes can yield the same field potential fluctuation (inverse model) (Buzsáki et al., 2012; Herreras, 2016; Herreras et al., 2015; Pesaran et al., 2018).

There is a general consensus that the prominent contributor to the low frequency component of the field potential (<100 Hz) is synaptic activity (Einevoll et al., 2007, 2013; Haider et al., 2016; Mazzoni et al., 2015; Mitzdorf, 1985), but interpretation remains complicated because the resultant field potential is significantly influenced by anatomical geometry, connectivity, tissue electrical properties, and nonsynaptic ionic currents (Buzsáki et al., 2012; Herreras, 2016; Lindén et al., 2010, 2011; Ness et al., 2016; Reimann et al., 2013). Though relatively simple models can capture certain relationships between the field potential and neuronal activity (Gao et al., 2017; Mazzoni et al., 2015; Miller et al., 2009), the details of the relationship between the two are mostly unknown.

Beyond the uncertainties of the specific current sources, the commonly used metrics of analysis (e.g., narrowband power) are often not specific and can be confounded with many different properties of the raw data, beyond that which is being conceptualized. For example, an increase in 10 Hz power can be a consequence of an increase in power of a: 1) stationary 10 Hz oscillator, 2) transient 10 Hz oscillator, 3) 5 Hz nonsinusoidal oscillator, 4) white noise, 5) a sharp transient, and more (Haller et al., 2018). Analyses of oscillations have mainly applied techniques based on the Fourier transform, which parametrizes signals as sums of sine waves at varying frequencies (Bruns, 2004; Cohen, 2017; Pesaran et al., 2018).

However, after sparse interest in the past (Amzica and Steriade, 1998; Jasper, 1948), recent interest has emerged regarding significance of the nonsinusoidal and nonstationary features of brain rhythms (Cole and Voytek, 2017; van Ede et al., 2018; Feingold et al., 2015; Fransen et al., 2015; Jas et al., 2017; Jones, 2016; Lozano-Soldevilla, 2018; Sweeney-Reed et al., 2018). Careful and thorough inspection of the neural signal, in the time-domain, is necessary in order to precisely characterize changes in neural oscillations and avoid potential pitfalls of spectral representations. By better parametrizing our features of interest, we are better suited to disentangle separate neural processes. To improve on conventional techniques, we have recently demonstrated a method of analyzing neural oscillations on a cycle-by-cycle basis, wherein we first determine when the oscillator is present, and then measure not only its power and frequency, but also its waveform symmetries (see Figure 1) (Cole and Voytek, 2018).

Parameterizing oscillation features on a cycle-by-cycle basis in this manner allows us to interrogate novel physiological relationships between neuronal spiking activity and multiple oscillatory features heretofore unexplored. It seems reasonable to assume that the properties of the oscillatory waveform correlate with the properties of the underlying physiological generators. There are several microcircuit motifs that produce oscillations (Womelsdorf et al., 2014), so changes of the specific motif or its components should manifest as changes in the waveform.

For example, three alpha rhythms in the gustatory cortex that each have distinct behavioral correlations can be differentiated by features of their waveforms (Tort et al., 2010). It is intuitive to interpret changes in oscillation amplitude and frequency as changes in the number of active neurons, or the time between consecutive activations, respectively (Pesaran et al., 2018). However, a conceptual relationship between the oscillatory waveform and neuronal activity is less apparent. Previously we hypothesized that the effect of deep brain stimulation treatment of Parkinson’s disease on making motor cortical beta oscillations more symmetric resulted from a desynchronization effect of the stimulation treatment (Cole et al., 2017). Given spike-field coupling, it may be expected that waveform symmetry would reflect the relative activity of different neuronal populations—e.g., excitatory and inhibitory ensembles or intra‐ and interlaminar interactions—given that the time windows during which neuron populations are active covaries with cycle symmetry.

Leveraging our novel cycle-by-cycle analysis approach, we sought to address several specific physiological hypotheses. First, we hypothesize that cycle-by-cycle variability in waveform shape can explain variance in neuronal firing rates, synchrony, and sequences. Next, we hypothesize that the presence of an oscillation, or burst, will change the statistics of local single-unit activity by increasing spiking, and spike synchrony, while stabilizing spike sequences. Finally, we hypothesize that the duration of an oscillatory burst can be predicted by the features of the very first oscillatory cycle in the burst.

To test these hypotheses, we analyzed a public data set of simultaneous recordings of the hippocampal local field potential (LFP) and spiking data from neuron units in region CA1 (Mizuseki et al., 2014). This data set was chosen because of the physiological properties of the hippocampal theta rhythm: its asymmetric waveform is stereotyped yet variable (Amemiya and Redish, 2018; Belluscio et al., 2012; Buzsáki et al., 1985; Trimper et al., 2014), there is established spike-field coupling with many hippocampal neuronal populations (Belluscio et al., 2012; Mizuseki et al., 2009, 2011), and the symmetry of the theta waveform has been linked to memory and representation (Amemiya and Redish, 2018; Trimper et al., 2014).

## Methods

Python code to replicate the figures in this paper are shared at https://github.com/voytekresearch/Cole_2018_theta.

### Data collection

Local field potentials (LFPs) and neuronal spiking were recorded from the CA1 pyramidal layer of the hippocampus in rats, and downloaded from the “hc3” dataset on the Collaborative Research in Computational Neuroscience (CRCNS) database (Mizuseki et al., 2014; Teeters et al., 2008). Briefly, extracellular recordings were made using multichannel silicon probes with 8 channels per shank (vertical distance: 20 μm) and either 4 or 8 shanks (200 μm spacing). Spikes were sorted at the original sampling rate (20 kHz or 32.552 kHz) and labeled as putative pyramidal neurons or putative interneurons based on their action potential waveforms and cross-correlations. LFP recordings were downsampled to 1250 Hz or 1252 Hz. Nine rats in this database had recordings from CA1. Recordings were downloaded from 3 sessions for each rat on 3 different days, if possible (27 total recordings sessions). Recordings were roughly chosen to maximize the number of simultaneously recorded pairs of interneurons.

The LFP recordings analyzed were taken from the deepest contacts on each shank. Normally, the traces from all recording contacts looked very similar due to their close proximity, but if the deepest contact significantly deviated from the other contacts, the next deepest contact was chosen. Recordings were collected from between 2 and 11 shanks in CA1 during each session. Shanks were analyzed independently (152 total shanks), and neurons were referenced to the theta recordings from the shank on which it was detected.

Five recordings analyzed also had simultaneous tracking data of the rat’s position. The speed during a theta cycle was computed as the distance traveled between the two peaks divided by the period. In order to analyze changes in hippocampal theta patterns during movement (Figure 3), theta cycles were conservatively classified as “moving” if the speed during that cycle was above the 90th percentile, and “stationary” if the speed was below the 10th percentile.

### Theta cycle analysis

The presence and features of hippocampal theta oscillations were analyzed using our previously described cycle-by-cycle analysis approach (Cole and Voytek, 2018). Briefly, a broad bandpass filter (1-25 Hz) was applied and then peaks and troughs were localized (Figure 1A, dots) in order to segment the signal into theta (4-10 Hz) cycles. Note this broad bandpass filter did not substantially affect the theta oscillation asymmetry of interest (compare gray and black traces in Figure 1A). A peak-to-peak segmentation was chosen because spiking was concentrated around the trough (Figure 4D,E) and so bursts of spiking around the trough would be analyzed in a single cycle (rather than 2 cycles if a trough-to-trough segmentation was used). For each cycle, four features were computed as shown in Figure 1B: amplitude, period, rise-decay symmetry, and peak-trough symmetry. Rise and decay midpoints were defined as the time points at which the voltage was halfway between the adjacent peak and trough voltages. These midpoints were used to represent the boundaries between peak and trough segments. Rise-decay symmetry is defined as the fraction of the period that is comprised of the rise phase. Peak-trough symmetry is similarly defined as the fraction of the period comprised of the peak phase, but the period in this case is bounded by consecutive rise midpoints instead of consecutive peaks.

It is important to appreciate that the neural oscillations are not present during the entire recording (Feingold et al., 2015; Jones, 2016; Lundqvist et al., 2016). Therefore, it is useful to determine the segments of the signal in which the oscillation is present because measuring theta features of a signal segment without a prominent theta oscillation will add noise to the analysis (Cole and Voytek, 2018). Therefore, only cycles that are determined to be part of a theta oscillatory burst were analyzed. However, the task of identifying the segments of the signal with oscillatory components is challenging and currently unsolved (Kosciessa et al., 2018). It is unclear if there are discrete times in which an oscillator is on and off, so perhaps there is no objective solution.

The approach for burst detection has been thoroughly described previously (Cole and Voytek, 2018), but briefly, a segment (cycle) of the signal was determined to be part of an oscillatory burst if its amplitude and period were comparable to adjacent cycles, and if its rise and decay flanks were mainly monotonic. Like with any burst detection algorithm, it relies on thresholds that must be semi-arbitrarily defined (Feingold et al., 2015; Hughes et al., 2012). In order to address this limitation, we ran our analysis with a range of burst detection parameters to assure that results were not simply dependent on one specific choice of settings. For the results shown in the main paper, the parameters were chosen as those that optimized the F1 score (equally weighted precision and recall) of a simulated signal with a signal-to-noise ratio that appears roughly similar to the hippocampal theta rhythm (Cole and Voytek, 2018). Thresholds were set such that adjacent cycles’ amplitudes and periods could be no more than 60% and 45% different, respectively, and the cycle flanks must be at least 80% monotonic. With these settings, theta oscillations were detected to be present 50-85% of the time across recordings.

### Neuronal spiking analysis

Spikes were previously sorted using KlustaKwik (Harris et al., 2000) and clustering was manually adjusted using autocorrelograms, cross-correlograms, and spike waveform shape (Mizuseki et al., 2014). Spikes were compared to the LFP recorded from the same shank. Neurons with fewer than 100 spikes during putative theta oscillations were excluded from analysis, resulting in a data set of 119 putative interneurons and 760 putative pyramidal neurons.

A measure of spike-field coupling (SFC) was computed for each neuron. The instantaneous phase of the LFP was estimated by interpolating between peaks (phase 0), troughs (π,−π), rise midpoints (−π/2), and decay midpoints (π/2), and the phase was determined at each spike time (Siapas et al., 2005). The distribution of these spike phases (50 circular bins, each of width π/25) was normalized by the distribution of phases in the recording in order to compute a firing rate in each phase bin. This normalization is critical because the consistent rise-decay asymmetry of the theta waveform results in more timepoints having positive phases compared to negative phases. SFC was then parametrized by the magnitude and phase of the mean vector, defined by summing each spike as a unit vector at the phase of firing. In the main manuscript, only spikes during theta bursts were included, as the theta phase is only reliable during these periods. In Supplementary Figure 1, these SFC estimates are compared to not having this theta burst requirement, as well as estimating the phase using the more conventional Hilbert transform-based approach, which has been shown to be systematically biased by waveform shape (Dvorak and Fenton, 2014).

In order to compare neuronal activity to hippocampal theta features (e.g., symmetry), firing rate was computed for each cycle by dividing the number of spikes in the cycle by the period. The correlation between firing rate and each theta feature was quantified by fitting a general linear model (GLM) to predict the firing rate from the cycle features. A one-sample Wilcoxon signed rank test was then used to assess if there is a significant bias in these model coefficients.

In supplementary analyses, we assessed if a cycle’s rise-decay symmetry was related to a neuron’s spike timing (Figure S2). However, there is no objectively correct reference frame by which to compute the time of a spike in a cycle, so three different ones were used in order to obtain a more holistic picture. For Figure S2A-D, the time for each cycle was normalized such that 0 corresponded to the former peak, and 1 corresponded to the latter peak. For Figure S2E-H, spike time was referenced to the trough of the cycle. For Figure S2I-L, phase was analyzed instead of spike time. In all cases, the modal spike time/phase was calculated for asymmetric and symmetric cycles, separately. Each cycle was classified as “asymmetric” (short rise, long decay) or “symmetric” using a threshold on the rise-decay symmetry (rdsym < 0.4: asymmetric; rdsym > 0.4, symmetric). A threshold of 0.4 rather than 0.5 was used because theta cycles were systematically more asymmetric with a shorter rise (rdsym < 0.4). To compute modal spike times, first spikes were binned (bin sizes: S2A-D: 5% of the cycle, S2E-H: 10 ms, S2I-L: π/10), and then this histogram was smoothed by convolving with a Gaussian (standard deviations: S2A-D: 5% of the cycle, S2E-H: 5ms, S2I-L: π/10). The spike time/phase with the highest firing rate (mode) was then determined for each neuron. This complicated method of binning and smoothing was designed to obtain a reasonable estimate of “average” since the mean and median do not have an intuitive interpretation when analyzing a circular process (periodic firing).

### Neuron synchrony and sequence analysis

Spike timing relationships between simultaneously recorded pairs of neurons were analyzed in order to test if the theta rhythm contained information about the firing patterns of the CA1 population. All simultaneously recorded pairs of neurons on the same shank were analyzed to identify events in which the two neurons fired within 20 ms of one another (a synchronous event). The neuron with fewer spikes was used as the reference neuron, and if the other neuron did not fire within 20 ms of a spike, that spike was labelled as a nonsynchronous event. If two synchronous events occurred in the same cycle, only one was maintained. If a synchronous and nonsynchronous event occurred in the same cycle, all events in that cycle were excluded. A neuron pair was analyzed if at least 100 synchronous events occurred during the theta rhythm (431 putative pyramidal neuron pairs, 46 putative interneuron pairs). The mean symmetries of cycles were calculated separately for synchronous and nonsynchronous events, and the difference was recorded for analysis across all neuron pairs (Figure 6A-D).

The fraction of spikes that were synchronous with the other neuron in the pair was computed separately during and not during theta bursts. The effect of the theta rhythm on synchrony was measured as the difference between these fractions (Figure 7F,G). This analysis of fractions necessitated sufficient samples in order to prevent one or a few events from significantly biasing results. Therefore, at least 25 synchronous and nonsynchronous events were required for both theta and non-theta periods (496 putative pyramidal neurons pairs, 45 putative interneurons pairs). Note that this is more neurons than were available for analysis with the aforementioned restriction of 100 synchronous events during the theta rhythm, because this requirement could be as few as 50 (25 pre + 25 post) synchronous events during the theta rhythm.

Synchronous events between neuron pairs (see above) were further analyzed to assess the neuron sequence. Synchronous events were defined as “pre” or “post” if the reference neuron fired before or after the other neuron. However, there are important cases in which a synchronous event should be excluded from synchrony analysis. For example, if the reference neuron fires a burst of 5 spikes, followed by a spike from the other neuron, this should not count as 5 “pre” sequences when doing statistics because these samples are not independent. In this case, in which multiple reference spikes occur within 20 ms, only the reference spike that is closest to the other neuron’s spike is kept for analysis. Additionally, If two reference spikes are recorded within 40 ms and are assigned opposite sequences, both events are removed from analysis. The ratio of “pre” and “post” sequences was computed, such that the ratio was always greater than 1, since the identity of the reference neuron was arbitrary. Thus, this “sequence ratio” corresponds to the relative bias of the neuron pair to fire among the two sequences. For example, a neuron pair in which neuron 1 fires before neuron 2 (1→2) 50 times and neuron 1 fires after neuron 2 (2→1) 75 times would have a sequence ratio of 1.5 (75:50 = 1.5:1).

## Results

### Cycle-by-cycle theta oscillation characterization

Field potential recordings from all shanks in CA1 were analyzed to characterize their hippocampal theta waveforms. A broad bandpass filter (1-25 Hz) was applied to the raw data in order to improve extrema localization while preserving the general shape of the theta rhythm (compare the gray and black traces in Figure 1A). After peaks, troughs, and flank midpoints were identified (see Methods), four features of each theta cycle were quantified (Figure 1B): 1) amplitude, 2) period, 3) rise-decay symmetry (“rdsym”), and 4) peak-trough symmetry (“ptsym”). Distributions of the cycle features are shown for an example recording. Note that the theta amplitude tended to be 2-3 mV (Figure 1C) and its period ranged between roughly 100 and 150 ms (7 - 10 Hz, Figure 1D). Notably, the theta cycles exhibited consistent symmetry biases. Specifically, the rise segment tended to be shorter than the decay (rdsym < 0.5, Figure 1E), and the peak segment tended to be shorter than the trough (ptsym < 0.5, Figure 1F).

We analyzed the consistency of these theta features across recordings. Recordings from most rats yielded cycle amplitudes around 2 mV, but recordings from 3 rats exhibited average amplitudes of 4-7 mV (Figure 1G). The average theta period was generally consistent across different recordings from the same rat (Figure 1H) and ranged roughly between 110 ms to 135ms (7.5-9 Hz). Rise-decay symmetry and peak-trough symmetry were below 0.5 in the vast majority of recordings (rdsym: 93%, ptsym: 95%), representing that the stereotyped theta waveform with short rises and short peaks is reliable across rats (Figure 1I-J).

**Figure 1.**
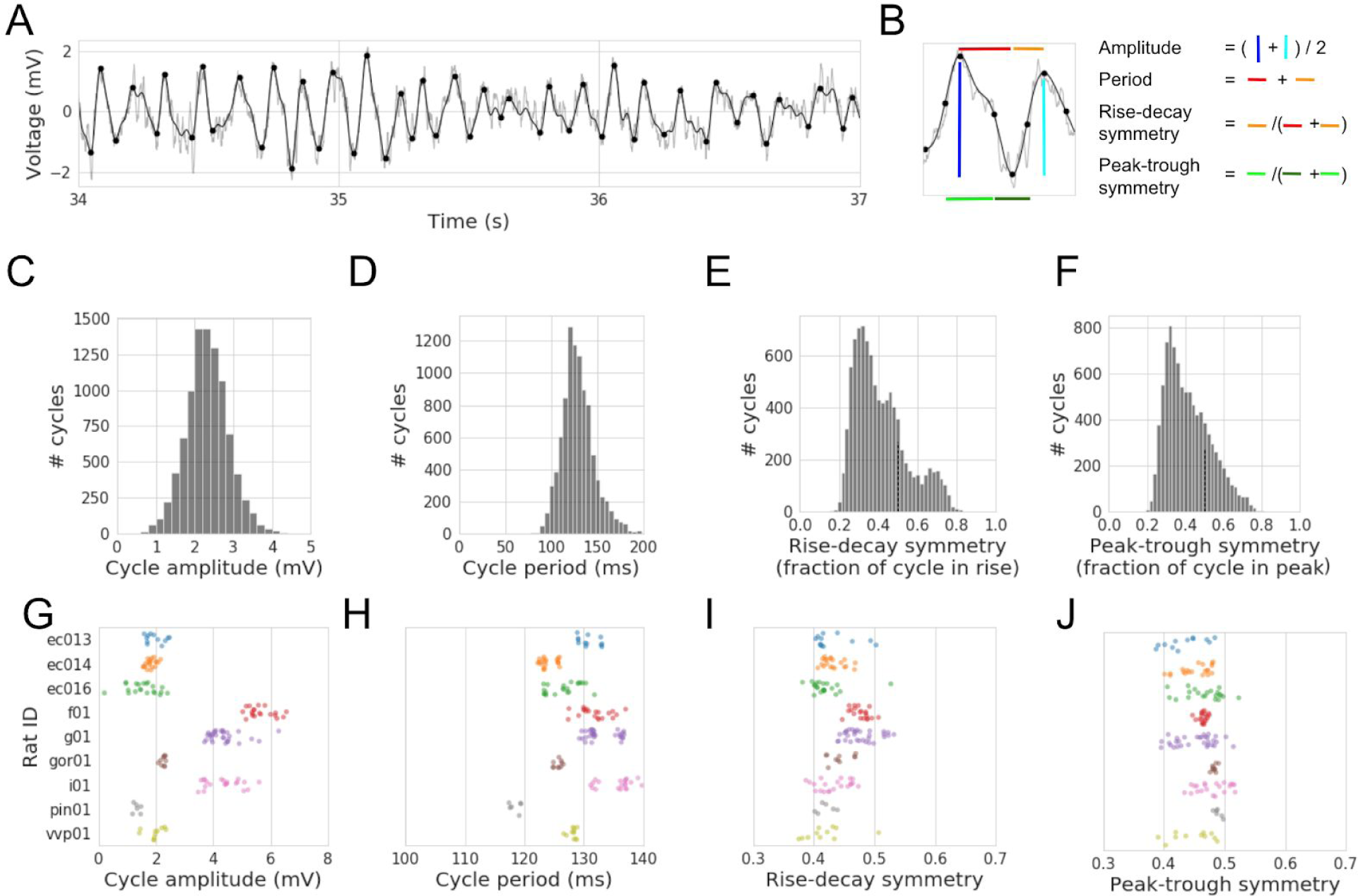
Cycle-by-cycle characterization of the rodent hippocampal theta rhythm. (A) Example trace of the local field potential recorded in the pyramidal layer of hippocampal CA1. The raw signal is plotted in light gray, and the black line shows the result of a broad bandpass filter (1-25 Hz). This broad bandpass filter reduces the high frequency noise that complicates extrema localization while still largely preserving the shape of the theta waveform. Identified peaks and troughs are denoted as black dots. (B) Illustration of how four features of the theta cycle are computed. Black dots in the middle of the rise and decay flanks denote the flank midpoints, which demarcate the boundary between peak and trough phases. Amplitude is computed by averaging the rise voltage (dark blue line) and decay voltage (light blue line). The period is the time between consecutive peaks (red and orange lines together). Rise-decay symmetry is defined as the fraction of the period in the rise phase (orange line). Peak-trough symmetry is defined as the relative length of time of the last peak (light green line) compared to the central trough (dark green line). (C-F) Distributions of (C) amplitude, (D) period, (E) rise-decay symmetry, and (F) peak-trough symmetry across all theta cycles in the example recording. Note that both symmetry measures are mostly below 0.5, indicating that these are non-sinusoidal, asymmetric oscillations wherein the rise tends to be shorter than the decay, and the peak tends to be shorter than the trough. (G-J) Distributions of the average (G) amplitude, (H) period, (I) rise-decay symmetry, and (J) peak-trough symmetry across all CA1 recordings in 9 different rats (each color). Note in (G) that 3 rats (f01, g01, and i01) have larger measured theta rhythms and in (H) that the recordings from the same rat tend to have consistent periods (i.e., steady theta frequency across recordings). Almost all recordings have, on average, relatively short (I) rise phases and (J) peak phases.

### Correlations between cycle features

When analyzing the significance of a single cycle feature, it is important to also account for the other cycle features if they are correlated with one another. Therefore, we characterized how the different cycle features correlated to one another. In Figures 2A-C, we explore in a single recording how each other cycle feature relates to rise-decay symmetry. Note that these distributions have significant structure that indicates mutual information (i.e., dependence) between these features. In order to identify a consistency in this structure, we summarized each pairwise relationship with a nonparametric Spearman correlation coefficient (ϱ) for each recording, and compared these ϱ values across recordings. We observed that theta oscillations that were more rise-decay asymmetric (shorter, or faster, rise) had a larger amplitude (Figure 2D, Wilcoxon signed rank test, N = 152, W = 2740, p < 10^−7^), shorter period (higher frequency, Figure 2E, W = 2835, p < 10^−7^), and had relatively longer peaks (Figure 2F, W = 28, p < 10^−25^).

Additionally, there were significant autocorrelations for each of the cycle features (Figure 2G). Theta amplitude (black line) is the most autocorrelated in the nearest cycles, followed by cycle period, and finally the cycle symmetries. Note that the dip in autocorrelation between periods of adjacent cycles reflects noise in peak localization, such that when a peak is detected artificially later, the latter cycle is artificially shorter and the former cycle is artificially longer. The relatively weak autocorrelations of rise-decay symmetry and peak-trough symmetry could reflect that these symmetry measures are inherently more noisy than estimates of amplitude and period and/or that oscillation asymmetry can better temporally resolve changes in physiology or behavior compared to amplitude and frequency.

**Figure 2.**
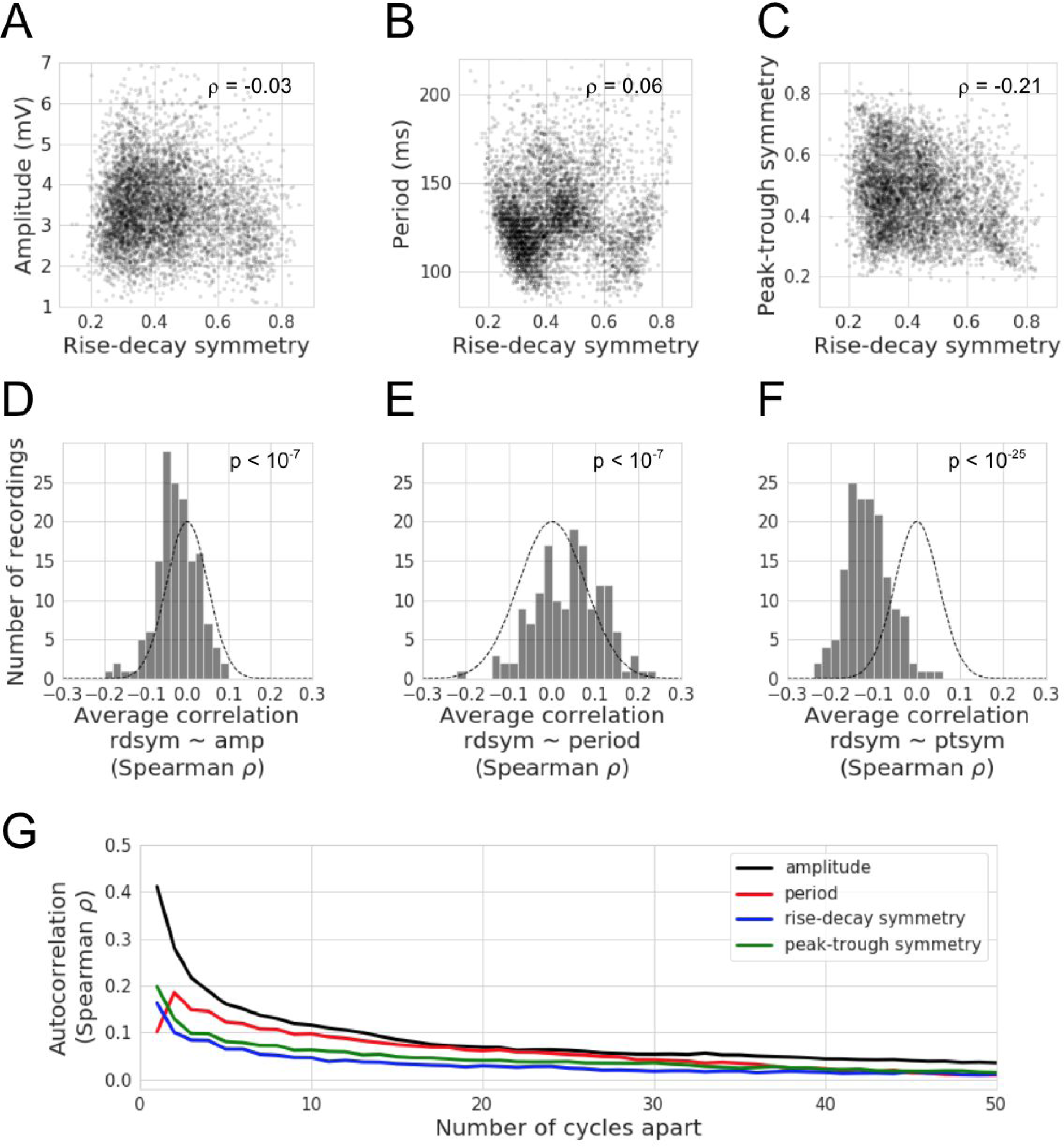
Correlations between features of hippocampal theta cycles. (A-C) For an example recording (same as in Figure 1A-F), there are significant correlations between the different cycle features. Each dot represents a single theta cycle. The rise-decay symmetry is slightly correlated to the cycle (A) amplitude, (B) period, and (C) peak-trough symmetry. (D-F) Distributions of Spearman correlation coefficients (ρ) across hippocampal recordings that relate the theta rise-decay symmetry on each cycle to its (D) amplitude, (E) period, and (F) peak-trough symmetry. Note that cycles that have a relatively short rise (rdsym < 0.5) tend to (D) have greater amplitude, (E) shorter periods, and (F) longer peaks (generally more peak-trough symmetric). Gaussian outlines are centered at zero with a variance equal to the distribution of the data, to visually compare results against the null hypothesis. (G) Lines show the correlation of features between cycles separated by increasing temporal distance (x-axis). The autocorrelations plotted are the averages across all 9 rats. Note that autocorrelations slowly decay over time, but remain consistently positive for several seconds (average cycle ~130ms). Amplitude (black) is the most autocorrelated, followed by the period (red), and the symmetries (blue: rise-decay, green: peak-trough) suggesting that there is more cycle-by-cycle independence of the symmetry metrics compared to amplitude or period.

### Rat movement and theta cycle features

For five sessions between two rats, spatial position data was available, and periods of fast movement and nonmovement were identified by computing the rat’s speed during each theta cycle (see Methods). Average theta cycle features were computed during these two types of periods, and significant differences were observed in all of them (Figure 3). Specifically, relative to nonmovement, while the rat was moving, the hippocampal theta oscillation was larger in amplitude (Figure 3A, N = 35, W = 78, p < 10^−3^), had a shorter period (faster frequency, Figure 3B, W = 0, p < 10^−6^), was more rise-decay asymmetric (Figure 3C, W = 4, p < 10^−6^), and was more peak-trough asymmetric (Figure 3D, W = 0, p < 10^−6^). Note these p-values should be interpreted with caution because the data come from only 2 different rats, so they do not necessarily generalize across the population. That said, these results are consistent with past reports of large, asymmetric, and relatively fast theta oscillations during running (Amemiya and Redish, 2018; Belluscio et al., 2012; Buzsáki et al., 1985; Hentschke et al., 2007).

In addition to these univariate statistics, we also fit a general linear model (GLM) to predict the rat’s speed during a theta cycle based on the four cycle features. This complementary approach is necessary because the cycle features are correlated to one another (Figure 2). Across sessions, the coefficients for period, rise-decay symmetry, and peak-trough symmetry were consistently negative, but the predictive sign of amplitude was not consistent (Figure 3E).

**Figure 3.**
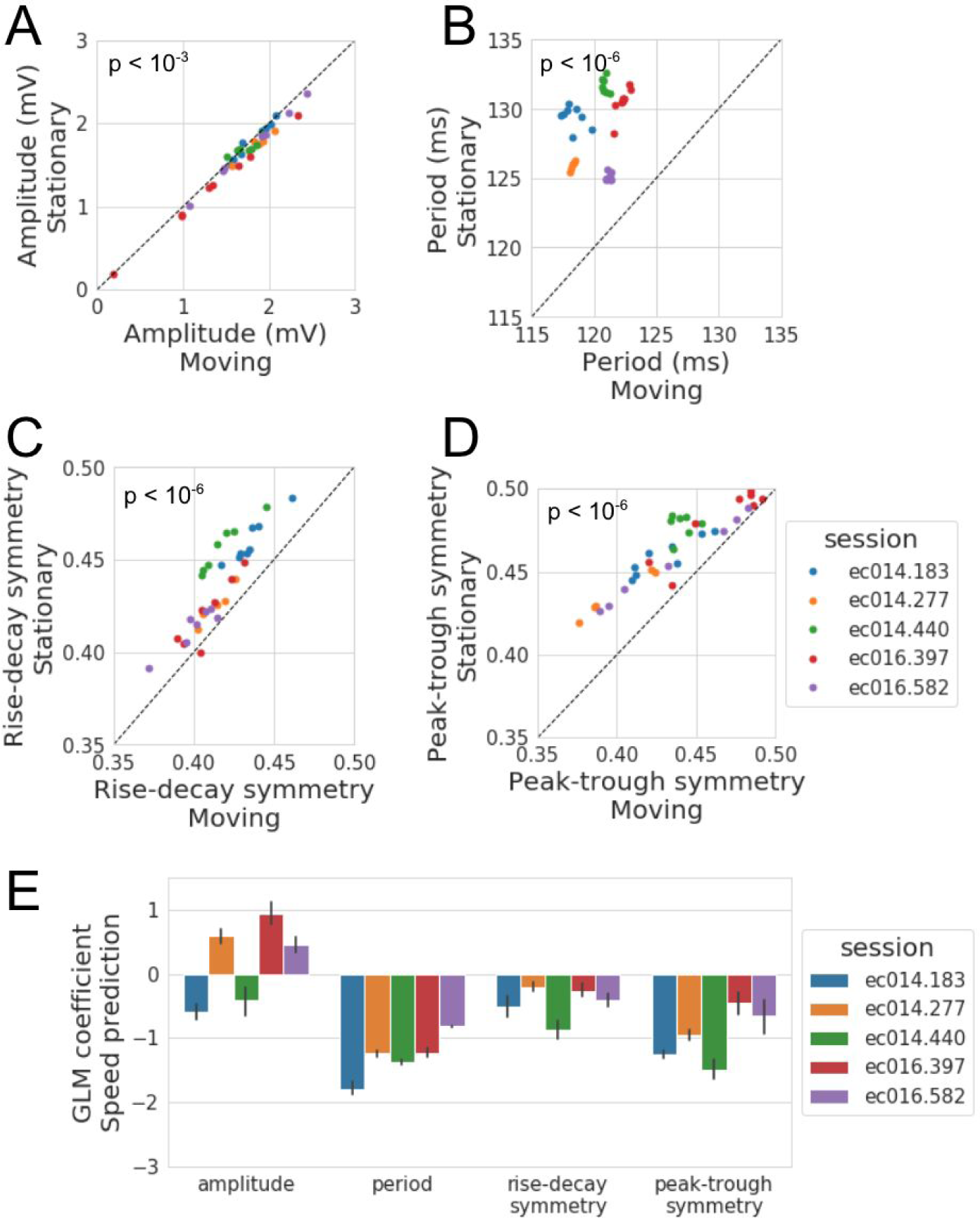
Comparison of hippocampal theta cycle features between movement and nonmovement. (A-D) Average properties of the decile of cycles in which the rat moved the most (x-axis, “Moving”) compared to the decile of cycles in which the rat moved the least (y-axis, “Stationary”). Each dot represents the recording from one shank in CA1, and each color represents a unique recording session for which position tracking was available from two rats. Note that in (A), the points mostly lie below unity (black line), indicating that theta amplitude is, on average, greater when the rat is moving. Additionally, while the rat is moving, (B) the theta period tends to be shorter, and (C) the cycles become both more rise-decay asymmetric (shorter rise) and (D) peak-trough asymmetric (shorter peak). (E) For each recording, a linear model was fit to predict the rat’s speed from the 4 cycle features. The bars show the average coefficient for each feature across all CA1 shanks simultaneously recorded in a session (error bars represent 95% confidence interval). Note that the amplitude feature does not consistently predict speed, but faster movement in all sessions was predicted by shorter periods and asymmetry values (i.e., more asymmetric with shorter rises and shorter peaks).

### Spike-field coupling between CA1 neurons and hippocampal theta

We now shift focus to how the features of the hippocampal theta rhythm relate to the network activity of neurons in hippocampal region CA1. Individual units were previously sorted and classified as putative pyramidal neurons and interneurons (Mizuseki et al., 2014) (see Methods). Figure 4A shows an example simultaneous recording of the theta oscillation (black) and spiking of a putative pyramidal neuron (red). As expected, the pyramidal neurons have significantly lower firing rates (Figure 4B, generally below 2 Hz) than the interneurons (Figure 4C, generally 20-40 Hz).

The correlation between neuron firing and the phase of the theta rhythm (i.e., spike-field coupling, SFC) has been well established in hippocampal neurons (Mizuseki et al., 2009, 2011). Figure 4A shows an example of this correlation for a pyramidal neuron with particularly strong SFC. This neuron consistently fires during the rise phase of the field potential (−π/2). Indeed, most pyramidal neurons fire at higher rates in the rise phase compared to the decay phase. Figure 4D shows the magnitude and phase of coupling as a black vector for each neuron, and the mean vector (red, 0.11*e*^−0.77π^) shows that the preferred phase for pyramidal neuron activity is in the rise period soon after the trough. In contrast, interneurons tend to fire in the decay phase prior to the trough (Figure 4E, mean vector 0.14*e*^0.85π^).

**Figure 4.**
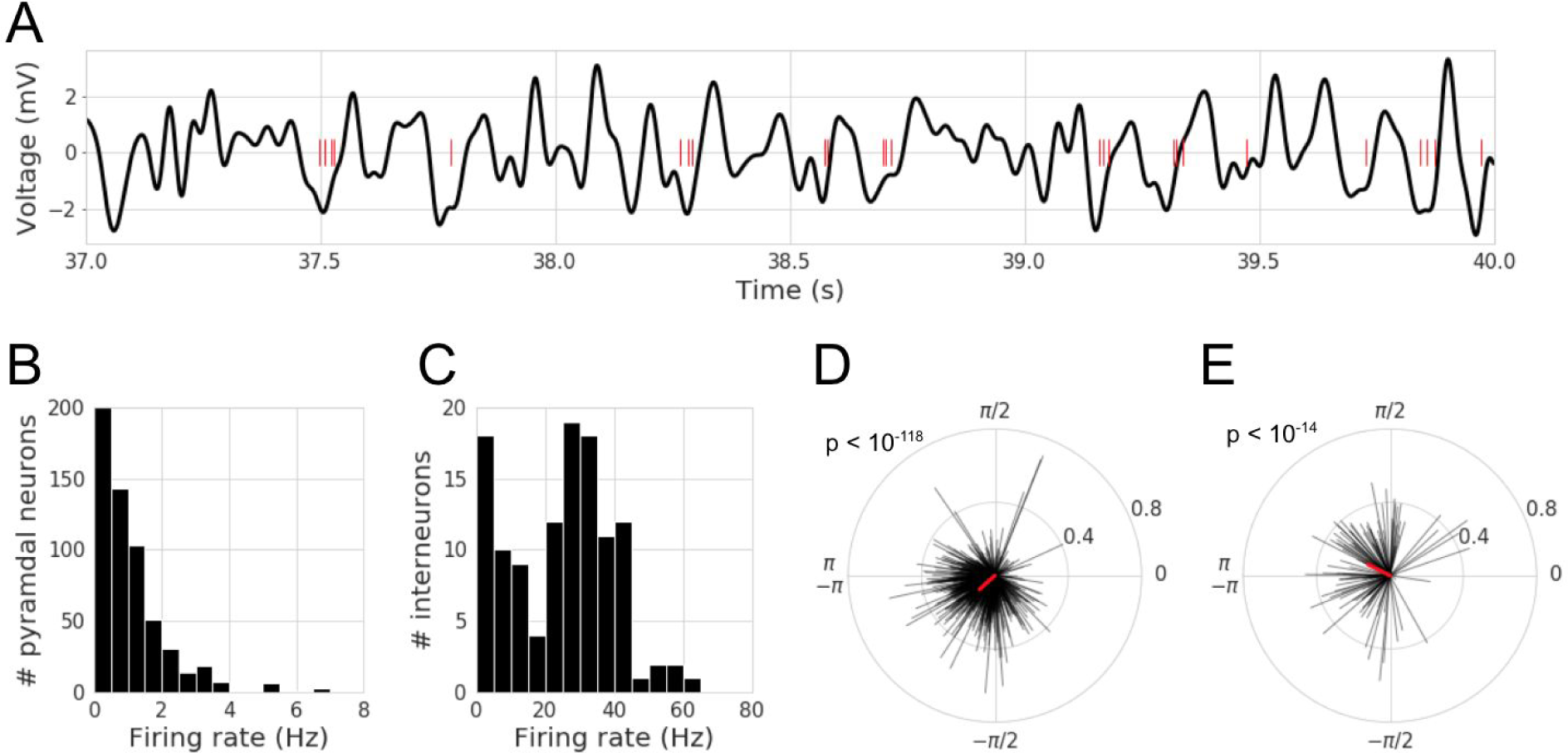
Spike-field coupling of CA1 neurons to the hippocampal theta rhythm. (A) Example CA1 field potential recording (black) and spike times (red) for a putative pyramidal neuron that tends to fire during the rise phase of theta oscillations. (B-C) Distributions of firing rates for all putative (B) pyramidal neurons, and (C) interneurons in the data set. Most pyramidal neurons have a firing rate below 2 Hz, and most interneurons fire between 20 and 40 Hz. (D-E) Distributions of spike-field coupling for putative (D) pyramidal neurons, and (E) interneurons. Each black line represents the coupling of a single neuron. The direction of the line reflects the phase at which the neuron is most likely to fire (phase 0 is the peak), and the magnitude of the line reflects the strength of the coupling. The red line shows the mean vector. Note that pyramidal neurons are most likely to fire after the trough (phase π/−π), while interneurons most likely fire before the trough.

In our analysis, we estimated SFC using the waveform-based phase estimate (see Methods) and only during periods of the signal in which theta was bursting. However, conventional SFC analysis uses portions of the signal in which the oscillation is not present, which negatively biases the coupling magnitude estimate (Supplementary Figure 1). Additionally, conventional approaches use a phase estimate based on the Hilbert Transform, which biases the phase estimate due to its narrowband filtering requirement and the nonsinusoidal nature of the theta rhythm (Supplementary Figure 1). In other words, burst detection and cycle-by-cycle parametrization enhances instantaneous phase and SFC estimates.

### Neuronal firing rate covaries with theta cycle features

As a complement to the well-known SFC effects, we analyzed the data in order to further identify relationships between neuronal firing and the LFP. GLMs were fit to predict the firing rate of neurons during each cycle from their normalized (z-scored) cycle features: amplitude, period, rise-decay symmetry (rdsym), and peak-trough symmetry (ptsym). The model coefficients (*β*) were recorded for each model and distributions of model coefficients across all neurons are shown in Figure 5A (pyramidal neurons) and 5B (interneurons). Both neuron types had higher firing rates during theta cycles with higher amplitude (pyramidal: N=760, W = 74416, p < 10^−30^, *β*_avg_ = 0.06, interneuron: W = 1072, p < 10^−10^, *β*_avg_ = 2.04), shorter periods (pyramidal: W = 66820, p < 10^−37^, *β*_avg_ = −0.08, interneuron: W = 337, p < 10^−16^, *β*_avg_ = −2.42), relatively short rise phases (rdsym, pyramidal: W = 117298, p < 10^−5^, *β*_avg_ = −0.04, interneuron: W = 613, p < 10^−14^, *β*_avg_ = −1.41), and relatively short peak phases (ptsym, W = 97066, p < 10^−14^, *β*_avg_ = −0.04, interneuron: W = 1366, p < 10^−8^, *β*_avg_ = −0.96). Not only is neuronal firing rate reflected by the commonly analyzed amplitude and frequency features, but it is also reflected significantly in the waveform symmetry.

**Figure 5.**
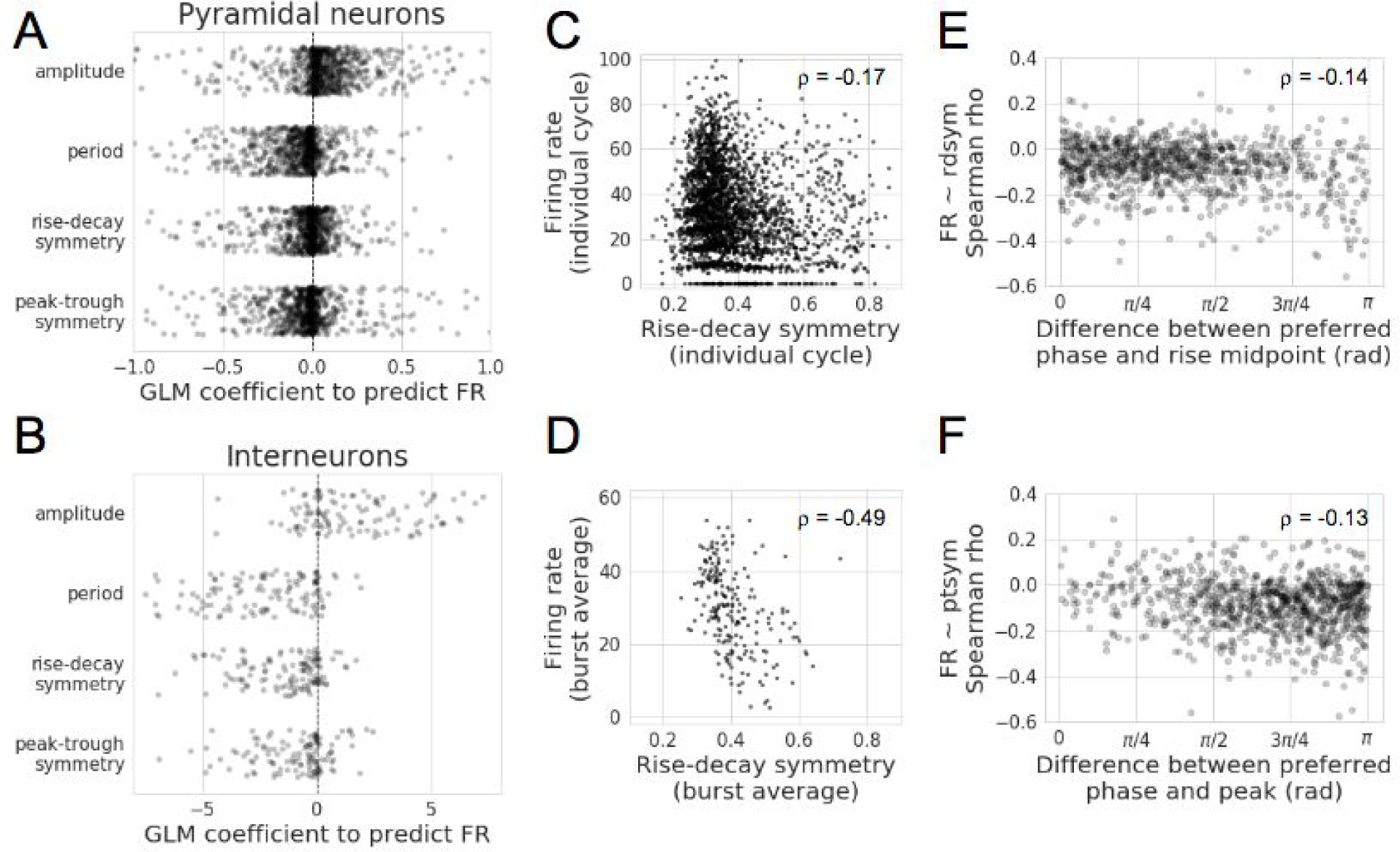
Theta cycle features are correlated to neuronal firing rate. (A-B) Linear models were fit to predict the firing rate of each neuron during a theta burst based on the average features of the component cycles. Each dot denotes the GLM coefficient for the model of the firing rate of an individual (A) pyramidal neuron or (B) interneuron. Increased theta amplitude and decreased theta period are associated with increased firing rates in both pyramidal neurons and interneurons. More rise-decay asymmetric (shorter rise) oscillations are also associated with faster firing rates in both pyramidal neurons and interneurons. More peak-trough asymmetric (shorter peak) oscillations are associated with faster firing rates in both pyramidal neurons and interneurons. (C-D) Visualization for a single putative interneuron that fires at a higher rate during cycles that are more rise-decay asymmetric (shorter rise), emphasizing the overall trend across all interneurons. Each dot represents one cycle in (C), or one burst in (D). Note that this correlation is even clearer when firing rate and rise-decay symmetry are averaged over a burst of cycles (D) compared to a single cycle (C). (E) The correlation between each neuron’s firing rate and rise-decay symmetry is compared to its preferred firing phase. Specifically, the x-axis shows the difference between the preferred firing phase and the rise midpoint (−π/2). Notice the negative correlation, indicating that the more that neurons prefer the decay phase, the more their firing rates tend to be stronger when the decay period is longer. (F) Similar to (E) but comparing the difference between a neuron’s preferred phase and the peak to the correlation coefficient between firing rate and peak-trough symmetry. This negative correlation indicates that if a neuron’s preferred phase is the trough, its firing rate is stronger when the trough period is longer.

We noticed that the cycle features that correlated to increased movement (larger, faster, more asymmetric) were in the same direction as those correlated to increased firing rate (Figure 3). Therefore, for the five sessions for which position tracking data was available, we tested if the symmetry features were still significant predictors of firing rate after the speed of the rat was accounted for. The GLM coefficients for these symmetry features remained consistently negative for both pyramidal neurons (N=760, rdsym: W = 19523, p < 10^−94^; ptsym: W = 14535, p < 10^−102^) and interneurons (N=119, rdsym: W = 50, p < 10^−19^; ptsym: W = 90, p < 10^−19^).

Note that the coefficient magnitude was largest for period and amplitude and smaller for the symmetry features, potentially indicating that the former features are generally more informative than the latter. Also note that the coefficients were an order of magnitude higher for the firing rate of interneurons, and the GLM’s average explained variance was 21% for interneurons compared to only 2.9% for pyramidal neurons. This difference can largely be attributed to the differences in firing rates between these neuron types, i.e., there will be high variance in the firing rate of a pyramidal neuron between theta bursts simply because these neurons fire more sparsely. This can be confirmed by noting a high correlation between a neuron’s firing rate and the GLM’s explained variance (Pearson r = 0.74).

When exploring univariate relationships, we noticed that effect sizes increased substantially if analysis was done with the basic unit of a burst as opposed to a single cycle. This is visualized for the relationship between firing rate and rise-decay symmetry for an example neuron. Rise-decay symmetry explains greater variance in firing rate when the fundamental unit is a burst (Figure 5D, ϱ =-0.49) compared to if the fundamental unit is a cycle (Figure 5C, ϱ =−0.17).

One potential explanation for a correlation between a neuron’s firing rate and theta asymmetry is that a neuron will fire at a higher rate if more time is spent in its preferred firing phase. In other words, if a single cycle of an oscillation has a very fast decay time, then each phase in that decay will last for less time than a more sinusoidal oscillation of the same frequency. Therefore, a neuron that prefers to fire during the decay phase will have a negative correlation between firing rate and rise decay symmetry (shorter rise, longer decay: increased firing). Therefore, we computed the circular-linear correlation (Berens, 2009) between a neuron’s preferred phase and this correlation coefficient and found that this distribution was significantly nonuniform (ϱ = 0.20, p < 10^−7^). Specifically, there was a significant negative correlation between this correlation coefficient (firing rate ~ rdsym) and the difference between a neuron’s preferred phase and the theoretical rise midpoint phase (−π/2) (Figure 5E, Spearman correlation, ϱ = −0.14, p < 10^−4^). These statistics support the aforementioned hypothesis that a neuron will fire at a higher rate if a longer part of the theta cycle is spent in its preferred phase.

This is extended by exploring the correlation between a neuron’s firing rate and theta peak-trough symmetry. Again, the circular-linear correlation found a nonuniform distribution in the neuron’s preferred phase and the correlation coefficient between firing rate and peak-trough symmetry (ϱ = 0.22, p < 10^−9^). Analogous to rise-decay symmetry, there was a significant negative correlation between this correlation coefficient (firing rate ~ ptsym) and the difference between a neuron’s preferred phase and the peak (Figure 5F, Spearman correlation, ϱ = −0.13, p < 10^−3^). Given these results, the nonsinusoidal waveform shape of the hippocampal theta rhythm seems to index the degree to which different neuronal populations are active, depending on their preferred firing phase.

### Theta oscillation features reflect neuronal synchrony and sequence

In addition to the firing rates of individual neurons, the coordinated activation of a population of neurons is thought to be important for neural computation. In particular, it is theorized that neurons transmit information more efficiently when they fire synchronously as opposed to asynchronously (König et al., 1996; Roy and Alloway, 2001; Salinas and Sejnowski, 2000; Stevens and Zador, 1998). We first attempted to analyze how differences in waveform symmetry related to differences in neuronal spike timing (Supplementary Figure 2), but this analysis proved to be prone to significant confounds (Supplementary Figure 3). Therefore, we investigated how features of the theta rhythm may correlate to synchrony between pairs of neurons. If these features differentiate degrees of synchrony, this would suggest that these oscillatory features contain important information about the function of the underlying neural oscillator.

Pairs of neurons were defined as firing synchronous when they fired within 20 ms of one another (see Methods). Compared to nonsynchronous spiking, during synchronous events, theta oscillations had increased amplitudes (pyramidal: N = 431 pairs, W = 24119, p < 10^−17^, 2.1% average amplitude increase; interneuron: N = 46 pairs, W = 179, p < 10^−4^, 4.2% average amplitude increase) and shorter periods (pyramidal: W = 30573, p < 10^−9^, 0.9% average period decrease; interneuron: W = 26, p < 10^−7^, 3.0% average period decrease). Additionally, synchronous interneuronal spiking tends to occur during both more rise-decay asymmetric (Figure 6A, W = 123, p < 10^−5^, 3.2% average decrease) and peak-trough asymmetric cycles (Figure 6B, W = 358, p = 0.046, 0.8% average decrease). The relationship between pyramidal neuron synchrony and asymmetry was weaker, if at all present (Figure 6C, rdsym: W = 43440, p = 0.23, 0.3% average decrease; Figure 6D, ptsym: W = 39459, p = 0.006, 0.5% average decrease). Together, these results show that theta cycle features contain some information about neuronal synchrony.

**Figure 6.**
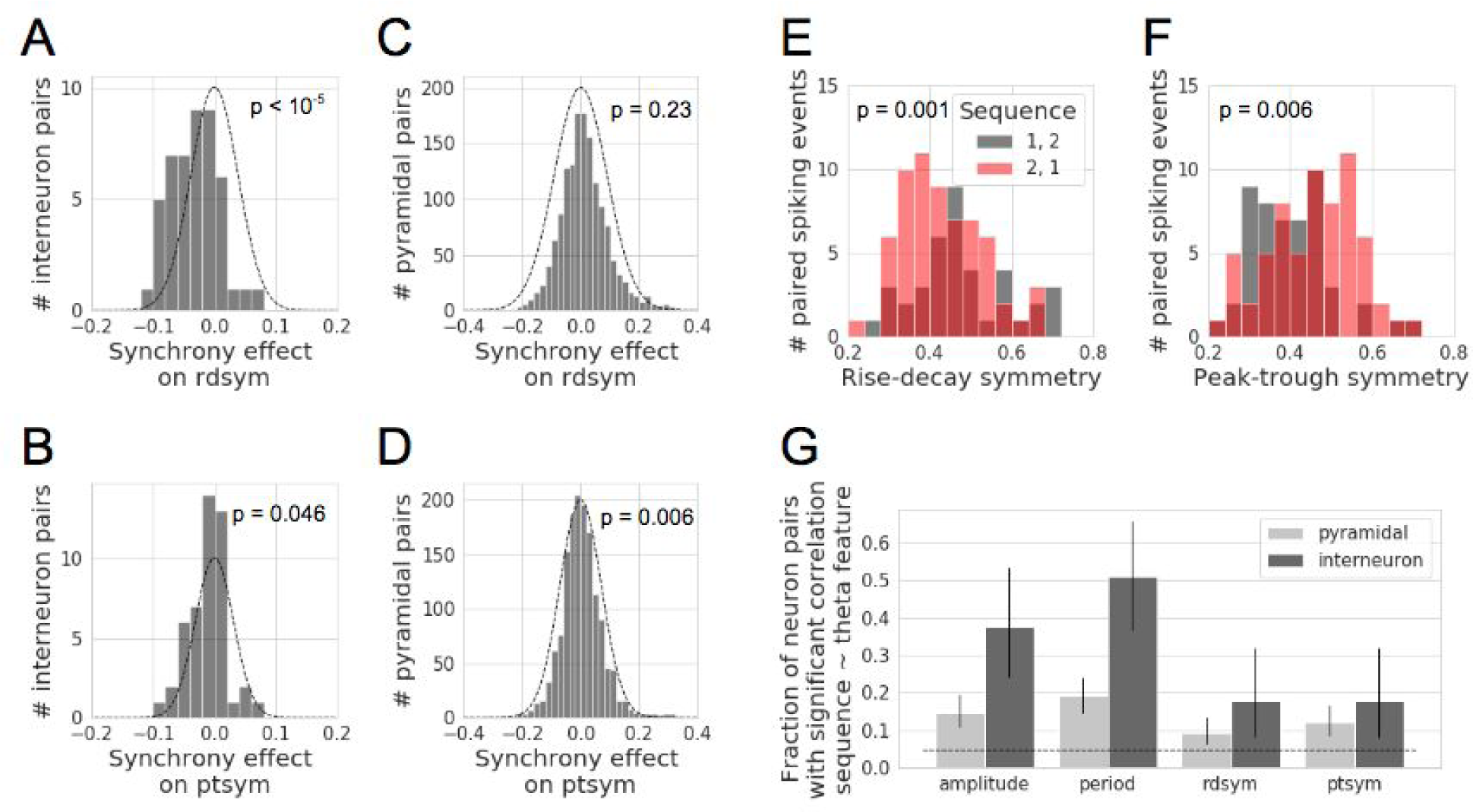
Neuronal synchrony and sequence are correlated with theta cycle features. (A-B) Distributions of the difference in (A) rise-decay symmetry and (B) peak-trough symmetry between synchronous events compared to nonsynchronous events for putative pairs of interneurons. Gaussian outlines are mean of 0 and *std* equal to the distribution of the data, to visually compare results against the null hypothesis. (C-D) Same as (A-B) but for pyramidal neurons, which do not have a strong relationship between their synchrony and the (C) rise-decay symmetry and (D) peak-trough symmetry of the ongoing hippocampal theta rhythm (E-F) Example pairs of putative pyramidal neurons showing significantly different distributions of (E) rise-decay symmetry or (F) peak-trough symmetry during cycles with one sequence compared to the opposite sequence. (G) Fraction of neuron pairs with a significant relation (One-sample Wilcoxon signed rank test, p < 0.05) between the sequence of firing and the theta cycle features. Note that the firing sequence of neuron pairs significantly relates to all four theta cycle features, and the effect is stronger for pairs of putative interneurons. Error bars denote the 95% binomial confidence interval for the number of significant neuron pairs. The dotted line denotes 5% of neurons that, by chance, would have a significant result.

We further investigated if synchrony between a simultaneously recorded pyramidal neuron and interneuron correlated to theta cycle features. We found similar trends as for pairs of interneurons such that pyramidal-inhibitory synchrony was related to higher amplitude, faster period, and more asymmetric theta cycles (N = 517 pairs; amplitude: W = 32508, p < 10^−23^, 2.7% average amplitude increase; period: W = 25841, p < 10^−32^, 1.8% average period decrease; rise-decay symmetry: W = 51641, p < 10^−5^, 0.8% average decrease; peak-trough symmetry: W = 47218, p < 10^−8^, 1.0% average decrease).

In addition to the importance of neuronal synchrony, the relative timing between neuron activations (sequence) is also theorized to reflect meaningful aspects of neural computation (Pastalkova et al., 2008; Skaggs and McNaughton, 1996; Wehr and Laurent, 1996; Yu and Margoliash, 1996). For example, neural circuit activation may be qualitatively different when neuron 1 (N1) fires before neuron 2 (N2) compared to when N1 fires after N2. Similarly, two oscillatory processes may be considered qualitatively different if in one, N1 and N2 have a regular sequence while in the second, N1 and N2 do not have a consistent firing sequence. Because the designation of N1 versus N2 is arbitrary, we measured sequence consistency as a “sequence ratio”, which was the ratio of synchronous instances in which N1 fired before N2 to when N1 fired after N2 (see Methods).

This allows us to analyze if the theta cycle features contained information about the underlying neuronal sequences. This could indicate that the symmetry of cycles may reflect differences in the state of the neuronal network and its computational roles. This would help explain a recent result in which neuronal activity could be used to decode current position better during more asymmetric (shorter rise) hippocampal theta cycles, and future position better during more symmetric cycles (Amemiya and Redish, 2018). In an example pair of neurons, the theta oscillation is mostly asymmetric (short rise) when N1 fires after N2, but more symmetric when N1 fires before N2 (Figure 6E, U = 950, p = 0.028). In another example neuron pair, the theta oscillation is more peak-trough asymmetric (short peak) when N1 fires before N2, but the oscillation is more symmetric when N1 fires after N2 (Figure 6F, U = 1098, p = 0.001).

We tested the significance of these sorts of effects across all eligible neuron pairs in our data set (see Methods). For each neuron pair, the distribution of cycle features was determined separately for the two sequences (N1 before N2, and N1 after N2), and a nonparametric, unpaired two-sample test (Mann-Whitney U) was applied to test if there was a significant difference in the cycle feature distribution between the two sequence events. The number of neuron pairs with a significant difference in cycle feature distribution (p < 0.05) was then compared to the number of neuron pairs expected by chance to have a significant effect (5%) using a binomial test (Figure 6G). It is notable that each cycle feature significantly correlated with firing sequence for both pyramidal neurons (amplitude: 41/278 pairs, p < 10^−9^, period: 53/278 pairs, p < 10^−16^, rdsym: 26/278 pairs, p = 0.002, ptsym: 34/278 pairs, p < 10^−5^) and interneurons (amplitude: 17/45 pairs, p < 10^−10^, period: 23/45 pairs, p < 10^−17^, rdsym: 8/45 pairs, p = 0.002, ptsym: 8/45 pairs, p = 0.002). Because spike order is arbitrary, we cannot overinterpret the results, though we can say there are consistent cycle feature differences for different spike sequences. These results were qualitatively similar when the time window of a “synchronous event” was varied between 10 ms and 50 ms. Furthermore, results were similar when investigating the sequence of neuron pairs consisting of one pyramidal neuron and one interneuron, and all tests withstood controls for multiple hypothesis testing by false discovery rate (FDR) correction (Benjamini and Hochberg, 1995).

### Relationship between theta bursting and cycle features

It is important to acknowledge that oscillations are not present in the signal at all points in time (Jones, 2016). This holds true for the hippocampal theta rhythm, perhaps the most stationary neural oscillation recorded in awake, behaving mammals, yet there are still periods in which it is clearly absent from the LFP. We used a burst detection algorithm (see Methods) to determine the time periods in which the theta oscillation was present. Figure 7A shows the distribution of burst duration throughout the data set. There was a minimum requirement of 3 cycles for an oscillatory period to be considered as a burst.

**Figure 7.**
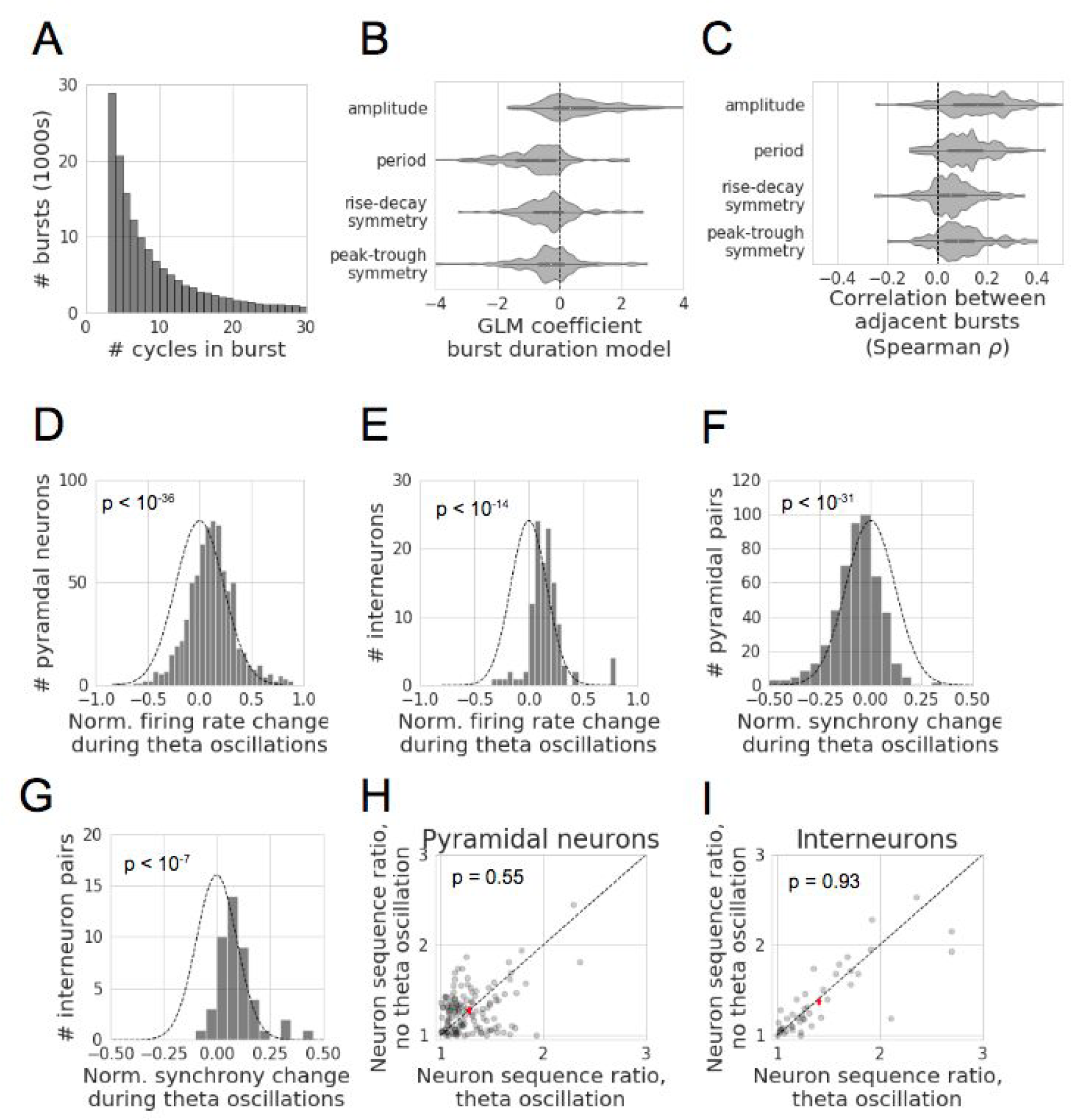
Characteristics of theta bursts and their relationship to cycle features and neuronal firing. (A) Distribution of durations of all theta bursts in the data set. The minimum burst duration was set to 3 cycles. (B) General linear models were fit to predict the duration of a theta burst based on the features of the first cycle. One model was computed for each hippocampal theta recording, and the distribution of coefficients across all recordings are shown. Note that theta bursts tend to be longer when cycles have higher amplitudes, shorter periods, and are more asymmetric (shorter rises and shorter peaks). (C) Correlation between average cycle features in adjacent bursts. Histograms show the distribution of Spearman correlation coefficients across all recordings. As indicated by distributions shifted to the right of zero, adjacent bursts tend to have more similar amplitudes, periods, and symmetries. (D-E) Distributions of the normalized difference in neuronal firing rate between periods of theta oscillation and no theta oscillation for putative (D) pyramidal neurons and (E) interneurons. Note that the distributions are shifted to the right of zero, reflecting that putative excitatory and inhibitory neurons fire more when a theta oscillation is present. Gaussian outlines are mean of 0 and *std* equal to the distribution of the data, to visually compare results against the null hypothesis. (F-G) Distributions of the normalized difference in neuronal synchrony between periods of theta oscillation and no theta oscillation for putative (F) pairs of pyramidal neurons or (G) pairs of interneurons. Note that during a theta oscillation, synchronous events between pyramidal neurons were less likely to occur (average 7% decrease), but more likely for interneuron pairs (average 9% increase). (H-I) Comparison of putative (H) pyramidal neuron and (I) interneuron sequence ratio during periods of theta oscillation and no theta oscillation. The “sequence ratio” measures the consistency in firing between a pair of neurons (neuron 1 and neuron 2, i.e., 1→2, or 2→1). Therefore, a sequence ratio of 1 represents both sequences occurred an equal number of times, and a sequence ratio of *x* means that one order was *x* times as common as the opposite order. Each dot represents a pair of neurons. The crosshare represents the mean and s.e.m. along each axis. Note for both neuron types, there is no significant difference in sequence ratio between periods with and without theta oscillations.

We used the burst detection method to test if there is a systematic (perhaps causal) relationship between the features of the first cycle in a burst and the burst duration. This could be the case if the properties of a specific physiological process that is related to oscillatory stability is detectable in the cycle features. For each recording, we fit a GLM to predict the burst duration from the amplitude, period, and symmetries of the first cycle. Features were normalized using a z-score relative to all cycles in that recording. We then assessed the consistency in the GLM coefficients by computing the coefficient distributions across recordings (Figure 7B).

Hippocampal theta bursts tended to be longer when the first cycle has a larger amplitude (N = 152, W = 2886, p < 10^−7^, average coefficient = 0.67), shorter period (W = 1262, p < 10^−16^, average coefficient = −0.78), is more rise-decay asymmetric (shorter rise, W = 2924, p < 10^−6^, average coefficient = −0.33), or is more peak-trough asymmetric (shorter peak, W = 3250, p < 10^−5^, average coefficient = −0.33). Therefore, all four cycle features are significantly predictive of the duration of the theta burst and together explain 4.9% of the variance in burst duration. Coefficients were qualitatively similar when speed was added as an additional predictor to the model for the neuronal firing rate during the sessions that contained position information (Figure 3). This suggests that these specific cycle features (high amplitude, short period, asymmetric) are indicative of a neural state in which the theta oscillation is more stable in time.

Segmenting the recordings into bursts also allows for comparing consecutive bursts. It is feasible that adjacent bursts would have similar features to one another, or for the cycle features to be relatively independent. The latter scenario would suggest that each burst of a theta oscillation is like an independent event that is unrelated to the previous burst of theta, which would have significant functional implications. To analyze this, we computed the average cycle features across all cycles in each burst. In most cases, we observe a positive correlation between the cycle features of adjacent bursts (Figure 7C). However, it is important to note that this is not always the case. There is not a significant correlation between adjacent bursts for average: amplitude in 22% of recordings, period for 30% of recordings, rise-decay symmetry for 47% of recordings, and peak-trough symmetry for 36% of recordings. Therefore, the dependence between adjacent theta bursts may depend on the specific context of the recording and local physiology.

### Theta oscillation presence relates to the firing rate, synchrony, and sequence of neuronal activity

Analysis of theta bursts on this dataset additionally allowed us to examine how CA1 neuronal network activity differed between periods of the recording with and without theta oscillations. For instance, both pyramidal neurons and inhibitory neurons tend to fire more during periods of the signal in which theta oscillations were detected (Figure 7D,E). This result was expected because of the established correlation between running and both increased firing rate and presence of theta activity (McNaughton et al. 1983). The presence of a theta oscillation is associated with an average 11% increase in pyramidal neurons (N=760, W=67682, p < 10^−36^), and 15% increase in interneuron, firing rates (N=119, W=569, p < 10^−14^). Again, we also studied synchrony between neurons. On average, synchronous events in a pair of pyramidal neurons had a 7% decreased likelihood during a theta oscillation burst (Figure 7F, N = 496 pairs, W = 23829, p < 10^−31^) despite the increase in pyramidal neuron firing rate during theta oscillations. In contrast, synchronous events were on average 9% more likely during a theta oscillation for pairs of interneurons (Figure 7G, N = 45 pairs, W = 43, p < 10^−7^).

We next analyzed if there was a difference in the relative consistency of neuron sequences between recording segments with and without theta oscillations. We hypothesized there to be a more consistent firing pattern during time periods with a prominent oscillation, with the idea that the oscillatory process is regularly firing a sequence of neurons. However, across all neuron pairs with a sufficient number of sequence observations (see Methods), there was no difference in the sequence consistency during segments with and without a theta oscillation (Figure 7H,I, pyramidal: N = 133, W = 4188, p = 0.55, interneuron: N = 44, W = 487, p = 0.93). This result was robust when varying the time window of sequence analysis between 10 ms and 50 ms.

## Discussion

This study provides a unique perspective on the rodent hippocampal theta oscillation by segmenting theta into individual cycles and analyzing how local spiking relates to nonsinusoidal cycle features and theta oscillation presence, as summarized in Figure 8. Such analyses are only possible using analytics based in the time domain, such as the cycle-by-cycle framework.

**Figure 8.**
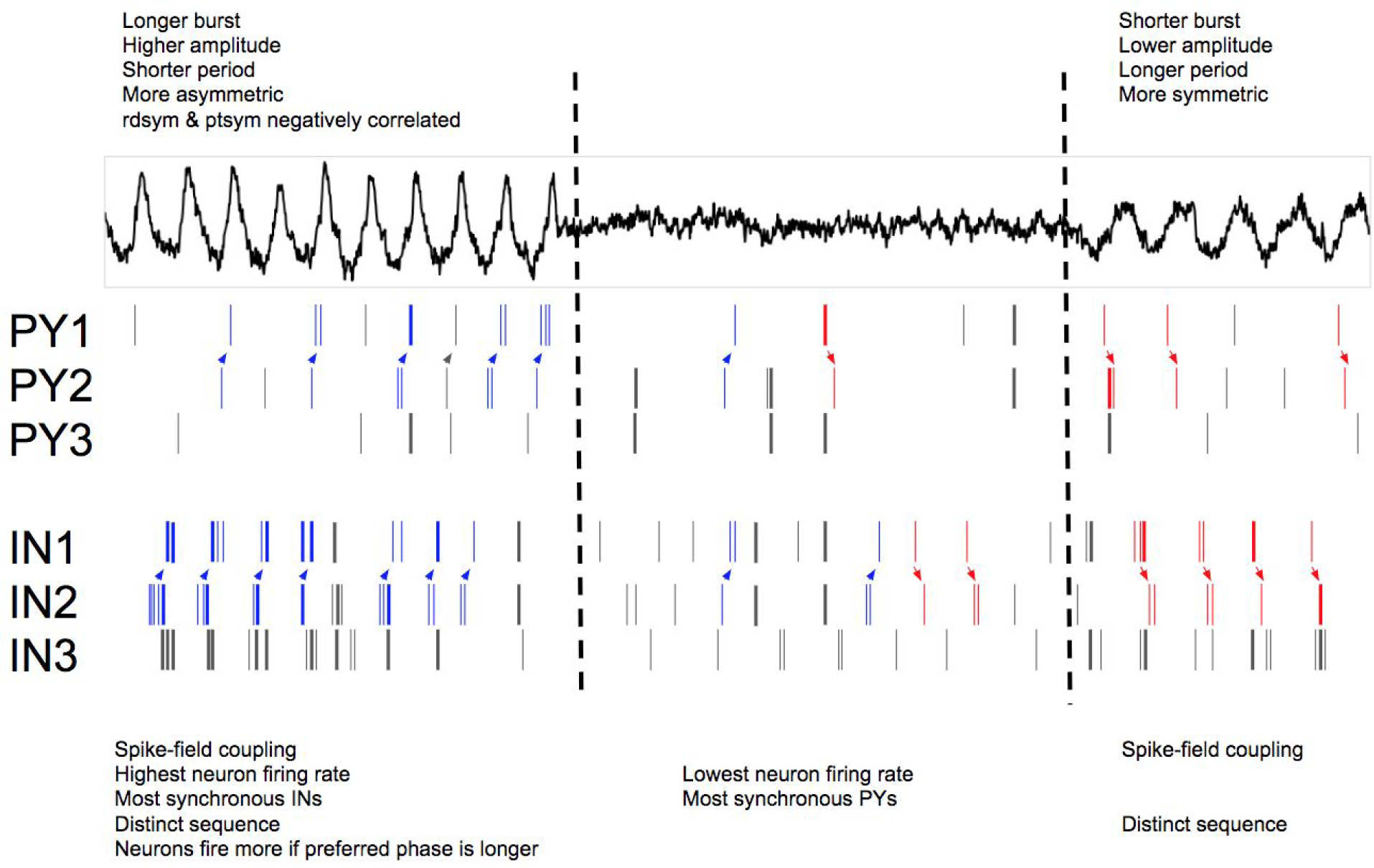
Schematized summary of observed relationships between theta oscillation bursting, waveform shape, and local neuronal firing patterns. A field potential (top) was simulated to show three different regimes of a hippocampal recording: a burst of asymmetric theta cycles (left), non-oscillatory activity (center), and a burst of symmetric theta waves. The asymmetric period would be most associated with the rat running, given its higher amplitudes, shorter periods, and relatively short rises and peaks compared to the latter symmetric theta burst (c.f. Figure 3). The rise-decay symmetry and peak-trough symmetry of asymmetric cycles are negatively correlated (c.f. Figure 2C,F). Additionally, the theta burst with more asymmetric cycles lasts longer than the symmetric burst (c.f. Figure 7B). Below the field potential is schematized firing of 3 pyramidal neurons (PY1, PY2, PY3) and 3 interneurons (IN1, IN2, IN3). Vertical lines indicate spikes from each neuron. The pyramidal neurons are coupled to the rise period after the trough, whereas the interneurons are coupled to the decay period before the trough (c.f. Figure 4). These neurons fire most when their preferred phase of firing is longest (c.f. Figure 5E,F) as well as firing most during the asymmetric burst (c.f. Figure 5, Figure 7D,E). During the asymmetric burst, interneurons are most synchronous (thick spikes) with one another (c.f. Figure 6B, Figure 7G). Pyramidal neurons are most synchronous when no theta rhythm is present (c.f. Figure 7F). The relative sequence of a pair of pyramidal neurons (PY1, PY2) and a pair of interneurons (IN1, IN2) are systematically different between the asymmetric and symmetric theta bursts (c.f. Figure 6E-G). Arrows are drawn and spikes are colored to clearly indicate the sequence of these neuron pairs (blue spikes are the sequence 2→1, red spikes are 1→2), though note that the sequences are arbitrary. However, when collapsing across both periods of theta oscillations, the sequence is no more stereotyped than during the time between theta bursts (c.f. Figure 7H,I).

Specifically, we uncover several novel characteristics of the nonsinusoidal theta oscillation and its relationship to putative excitatory and inhibitory firing rates, spike synchrony, and spike sequences. For one, the peak-trough asymmetry of the theta rhythm has not previously been parameterized or studied, but here we find that it has a characteristic asymmetry that systematically relates to both rat movement and neuron sequence. Our analysis of rise-decay symmetry extends significantly on the previous, mostly qualitative, reports of the sawtooth-like nature of the hippocampal theta rhythm (Amemiya and Redish, 2018; Belluscio et al., 2012; Buzsáki et al., 1985; Hentschke et al., 2007).

Recent work has shown that the theta rise-decay symmetry correlates with the ability to decode present or future position from place cell firing (Amemiya and Redish, 2018). Specifically, current position was more accurately decoded during asymmetric theta cycles, and future position was better decoded during symmetric cycles. Here we extend this result by demonstrating that this rise-decay symmetry correlates with local pyramidal neuron and interneuron firing patterns. Therefore, the higher firing rates, greater synchrony, and specific pairwise sequences in CA1 during asymmetric cycles may be important neural computational elements for representing current position, while representing future position is supported by lower firing rates and synchrony and alternative neuronal sequences.

### Non-independence between cycle features and adjacent cycles

During analysis, awareness that the cycle features are not independent of one another (Figure 2A-F) is critical. Nonsinusoidal waveform shape is complex, with many possible features for parametrization. Additionally, it is important to recognize that features are not independent across cycles (Figure 2G). The observations that all cycle features are significantly correlated, not only in adjacent cycles but also with nearby cycles, indicates the speed at which the oscillatory dynamics can change. Concretely, neural activity is more similar in two theta cycles within a few seconds compared to two theta cycles that are minutes apart. Because of this lack of independence across cycles, the p-values for statistical tests within a recording should be cautiously interpreted. Therefore, in this paper, we instead computed a single statistic for each recording (e.g., correlation coefficient between firing rate and rise-decay symmetry) and tested if the statistics are randomly distributed around zero.

### Burst detection and analysis of oscillatory time periods

Algorithmic determination of theta burst periods allowed us to study how cycle features vary across bursts of theta oscillations. Across cycles, we observed that the theta features were consistently autocorrelated (Figure 2G). It is possible this autocorrelation does not extend across distinct theta bursts but only exists within a burst. However, Figure 7C showed that the average cycle features of adjacent bursts are generally positively correlated. This could allow us to conclude that the neural dynamics that periodically occur during theta cycles slowly change over time. However, as mentioned in *Methods*, burst detection algorithms are not perfect, so this result should not be considered definitive. Autocorrelation of cycle features between bursts could potentially be an artifact of the algorithm artificially splitting a single burst into multiple bursts due to noise from aperiodic components of neural activity. In this scenario, the truth may be that when a theta burst ends and a new one begins, their oscillatory dynamics are essentially independent of one another. This interpretation still is possible for the minority of the recordings in which no correlation was observed between features in adjacent bursts (Figure 7C).

This prediction of burst duration based on cycle features (Figure 7B) reflects an interesting aspect of the underlying physiology, in which oscillatory network dynamics that produce more asymmetric field potentials are more stable than those that produce more symmetric waveforms. This is reminiscent of previous modeling studies that showed that more asymmetric oscillators synchronize more quickly than more sinusoidal oscillators (Somers and Kopell, 1993). Note that theta is generally faster and more asymmetric during running periods (Figure 3), and it is generally known that hippocampal theta is persistent during running periods (McFarland et al., 1975; Teitelbaum and McFarland, 1971; Vannderwolf, 1964). Therefore, this observed correlation between cycle features and stationarity is likely partly a consequence of analyzing periods of movement and non-movement together. That said, as stated in the Results, these trends held when accounting for speed in the linear model to predict burst duration from the cycle features. Future analysis using data with thorough behavioral tracking and annotation can tease apart this result in more detail.

### Relationship between theta oscillation and neuronal activity

In addition to analyzing trends within the LFP, a burst detection algorithm also opens the possibility for comparing neuronal activity between periods with and without a theta oscillation. We observed that, in general, both putative pyramidal neurons and interneurons increase in firing rate during a theta oscillation (Figure 7A,B). From this result, it would be expected that there would be more synchrony between neurons (i.e., pairs of neurons would more likely fire together within a short time window). Indeed, this was observed for interneurons (Figure 7B), but surprisingly the opposite trend was observed in general for pyramidal neurons (Figure 7A). That is, pyramidal neurons were less synchronous with one another during theta oscillations. From this, we conclude that theta oscillations do no necessarily enhance synchrony between neurons in a local population, which may not be expected given the widespread idea that oscillations synchronize neurons (Engel et al., 1990, 1997, 2001; Fries, 2005; Livingstone, 1996; Singer, 1999). This idea should be further explored regarding the relative synchronization of neurons between regions, or in higher frequency (beta and gamma) bands.

We find that neurons fire at higher rates if a longer part of the theta cycle is spent in its preferred phase. That is, when an individual theta cycle, for example, rises too quickly, a neuron that prefers a rise phase will have less time to fire at its preferred phase because the oscillation cycled too quickly out of that preferred phase window. However, if that cycle rises more slowly, the neuron will have a longer temporal window in which to fire, as that cycle spends more time in its preferred phase. One potential consequence of this observation is that asymmetries in individual cycles may be a means for controlling representation and/or computation, wherein specific subnetworks of neurons with different preferred phases can be selected for by specific cycle asymmetries. That is, this introduces the notion of neuronal networks that are controlled by the temporal duration of nonsinusoidal waveform features.

Similarly, there is also the idea that during a neural oscillation, neurons fire in a particular sequence that is not present in the absence of the oscillation (Mehta et al., 2002; Roux and Uhlhaas, 2014; Wehr and Laurent, 1996). However, we find no evidence of this in the hippocampal theta rhythm, as the firing sequence in neuron pairs was no more consistent during theta oscillations compared to periods without theta oscillations (Figure E-F). This may indicate that neuronal sequences are independent of the presence of the theta rhythm. It is still theoretically possible that higher order sequences (3+ neurons) are more consistent during theta oscillations, but more complex methods are required to assess this scenario. Even then, it is rather strange to consider a scenario in which theta oscillations coordinate longer sequences while having no effect on pairwise sequence consistency.

### Future directions

This manuscript serves as the first work to systematically relate nonsinusoidal waveform features of a neural oscillation to local neuronal spiking patterns. We showed that neuronal activation patterns correlate with the waveform shape of the theta oscillation, suggesting that changes in the symmetry of an oscillation may reflect qualitative changes in the function of the oscillation. This established relationship to spiking may help motivate and inform applications of nonsinusoidal features as predictive features for brain-machine interfaces or biomarkers for disease.

This work can be extended in several ways to further explore the potential significance and interpretations of oscillatory waveform asymmetries. It may be advantageous to study spatial patterns of the field potential (Agarwal et al., 2014). Additionally, studying simultaneous intracellular voltage and extracellular recordings may elucidate how fluctuations in the field potential relate to excitatory and inhibitory synaptic input. Since the field potential is not generated simply from the local spiking patterns (Herreras, 2016) neuronal activity from different brain regions may account for significant variance in the field potential recordings.

In addition to the rodent hippocampal theta rhythm, the relationship between the LFP waveform and local spiking patterns should be conducted in different frequency bands, brain regions, and species. Furthermore, new techniques may be developed to capture important features of oscillatory dynamics. For example, the “smoothness” of the oscillation was not studied here, and state-of-the-art deep learning methods could provide some informative, though less intuitive, features. Ultimately, this work only scratches the surface of future efforts to better understand the physiological, behavioral, and functional significance of neural oscillation waveform shape.

## Supplementary Text

### Different strategies of estimating spike-field coupling of CA1 neurons to hippocampal theta

One goal in analyzing neural oscillations is towards determining how they reflect underlying physiological changes. The “spike-field coupling” phenomenon in which neurons are more likely to fire at a particular oscillatory phase has been previously reported in the theta rhythm (Mizuseki et al., 2009, 2011). Because phase is commonly computed using a sinusoidal decomposition, or by relying on a filter parametrized using the Fourier Transform, previous studies have used the theta waveform to obtain a more accurate estimate of instantaneous phase (Belluscio et al., 2012; Siapas et al., 2005).

This choice is important, as we show that the coupling estimate is biased by the chosen method (Figure S1A-B). For pyramidal neurons coupled to the end of the rise phase, applying the conventional technique for phase estimation based on the Hilbert transform (Le Van Quyen et al., 2001; Lee et al., 2005) results in the estimated coupling magnitude to be relatively higher than the waveform-based phase estimate (Figure S1C, Spearman correlation, π = 0.48, p < 10^−137^). However, this Hilbert transform method relatively underestimated coupling for neurons coupled to the beginning of the rise phase. For interneurons, the Hilbert transform method gives a higher estimated coupling magnitude during the rise phase and a lower estimated coupling magnitude for neurons coupled to the decay phase (Figure S1D, π = 0.56, p < 10^−39^). Additionally, the coupling phase is biased by the chosen estimation technique. Figure S1E shows that the estimated preferred phase in pyramidal neurons is later when using the Hilbert transform technique compared to the waveform-based phase estimate (W = 795532, p < 10^−81^). For interneurons, the preferred phase estimate difference is phase-dependent (Figure S1F). These trends are likely due to the artificial symmetry enforced onto the asymmetric theta waveform when a theta bandpass filter (4-10 Hz) is applied. We further show that the coupling magnitude is underestimated when a burst detection algorithm is not used (Figure S1G-I, W = 1144140, p < 10^−111^).

**Figure S1.**
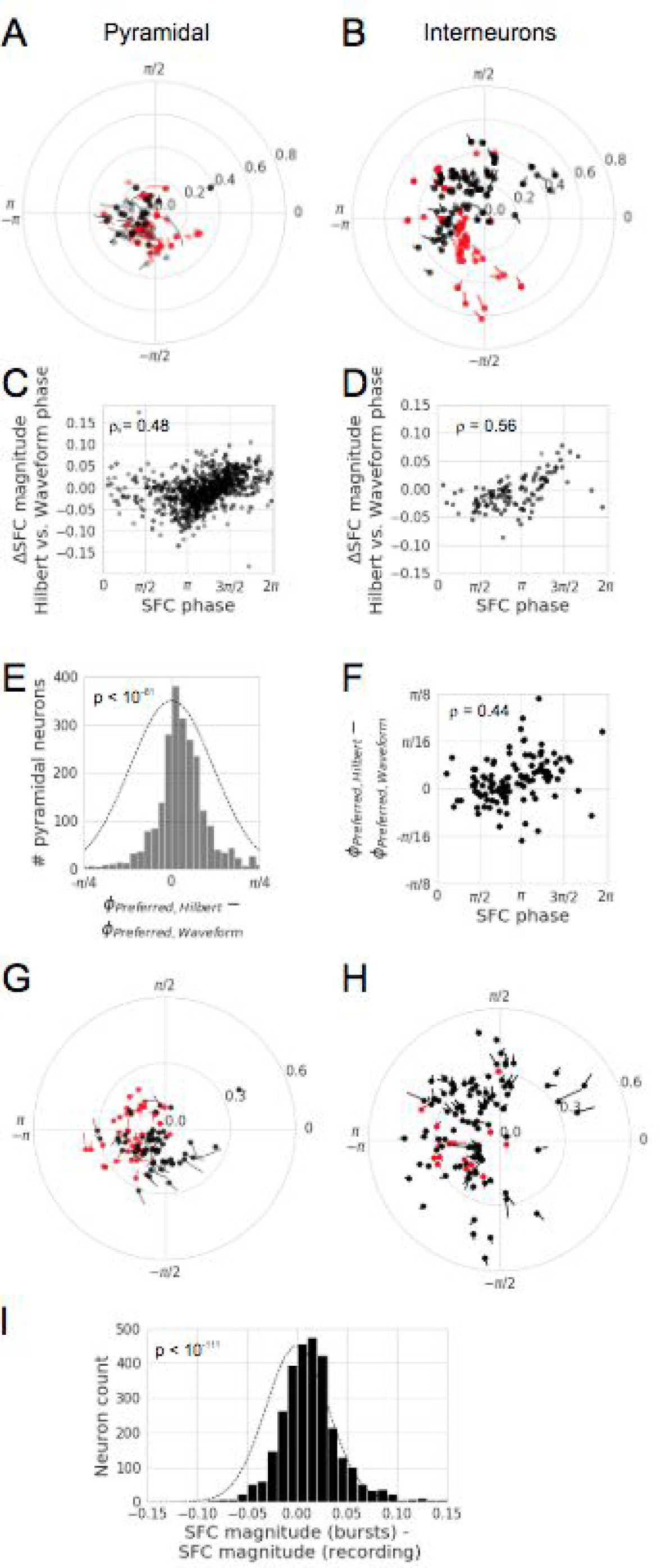
Differences in spike-field coupling estimates between techniques. (A-B) Differences in spike-field coupling estimates when using the conventional Hilbert transform phase estimate (circles) versus the cycle-by-cycle waveform estimate (ends of line). A sample of 100 pyramidal neurons are shown in (A), and all putative interneurons are plotted in (B). Red lines denote comparatively higher SFC magnitude when using the Hilbert phase estimate. A lighter dot represents a later preferred phase estimate when using the Hilbert transform technique. (C-D) SFC magnitude estimates differ between the Hilbert transform and waveform estimates depending on the phase of coupling. The x-axis corresponds to the phase estimate using the waveform approach. For putative pyramidal neurons (C) coupled to the trough (−π/π), the estimate of SFC magnitude is higher using the waveform phase estimate compared to the Hilbert estimate. However, the SFC magnitude is comparatively lower using the waveform estimate when the neuron is coupled to the theta peak. For interneurons (D) coupled to the rise period (-π, 0), the SFC magnitude is higher using the waveform phase estimate, but the estimated SFC magnitude is comparatively lower for neurons coupled to the decay period (0, π). (E) Difference between estimates of putative pyramidal neuronal preferred firing phase. The Hilbert transform-based estimate results in a preferred phase that is consistently later in the cycle than the waveform phase estimate. The Gaussian outline is centered at zero with a variance equal to the distribution of the data, to visually compare results against the null hypothesis. (F) For putative interneurons, the difference between preferred phase estimates is heavily correlated with the preferred phase. During the decay period, the Hilbert transform-based preferred phase estimate is comparatively earlier in the cycle, but if the neuron is coupled to the rise period, the Hilbert transform-based preferred phase estimate is comparatively later in the cycle. (G-H) Spike-field coupling estimate difference between using the whole time series (circles) or only the cycles that were defined as during a theta oscillation (end of line). A sample of 100 putative pyramidal neurons are shown in (G), and all interneurons are plotted in (H). Note that in most cases (black dots and lines) that the estimate of SFC magnitude is lower when the whole signal is used for estimating spike-field coupling, rather than only the time periods identified as theta oscillations. Otherwise, the samples are colored red. (I) The distribution of SFC magnitude differences shown in G & H showing a consistently higher SFC magnitude when computed only during theta bursts compared to computed over the whole recording.

### Assessing changes in spike timing with rise-decay symmetry changes

The first way that we attempted to test if theta symmetry reflects a change in the local network activity was by looking at changes in spike timing. The idea was that, if theta cycle rise-decay symmetry reflects differences in neural circuit activation, then neurons may fire at relatively different times within the cycle during asymmetric compared to symmetric cycles. First, we normalized time such that 0 corresponded to the former peak and 1 corresponded to the latter peak and analyzed these distributions of spike times. We observed that both pyramidal neurons and interneurons fired at earlier normalized times in the cycle (Figure S2A-D). However, the phase progression over normalized time systematically differs between asymmetric and symmetric cycles, which could lead to a confounding result. Therefore, we also studied changes in spike timing with two other reference schemes. When time-locking to the trough of the cycle, spikes occurred later during asymmetric cycles (Figure S2E-H). This was also the observation when spike phase was used (Figure S2I-L). Together, these data suggested a complex change in spike timing as a function of cycle symmetry.

**Figure S2.**
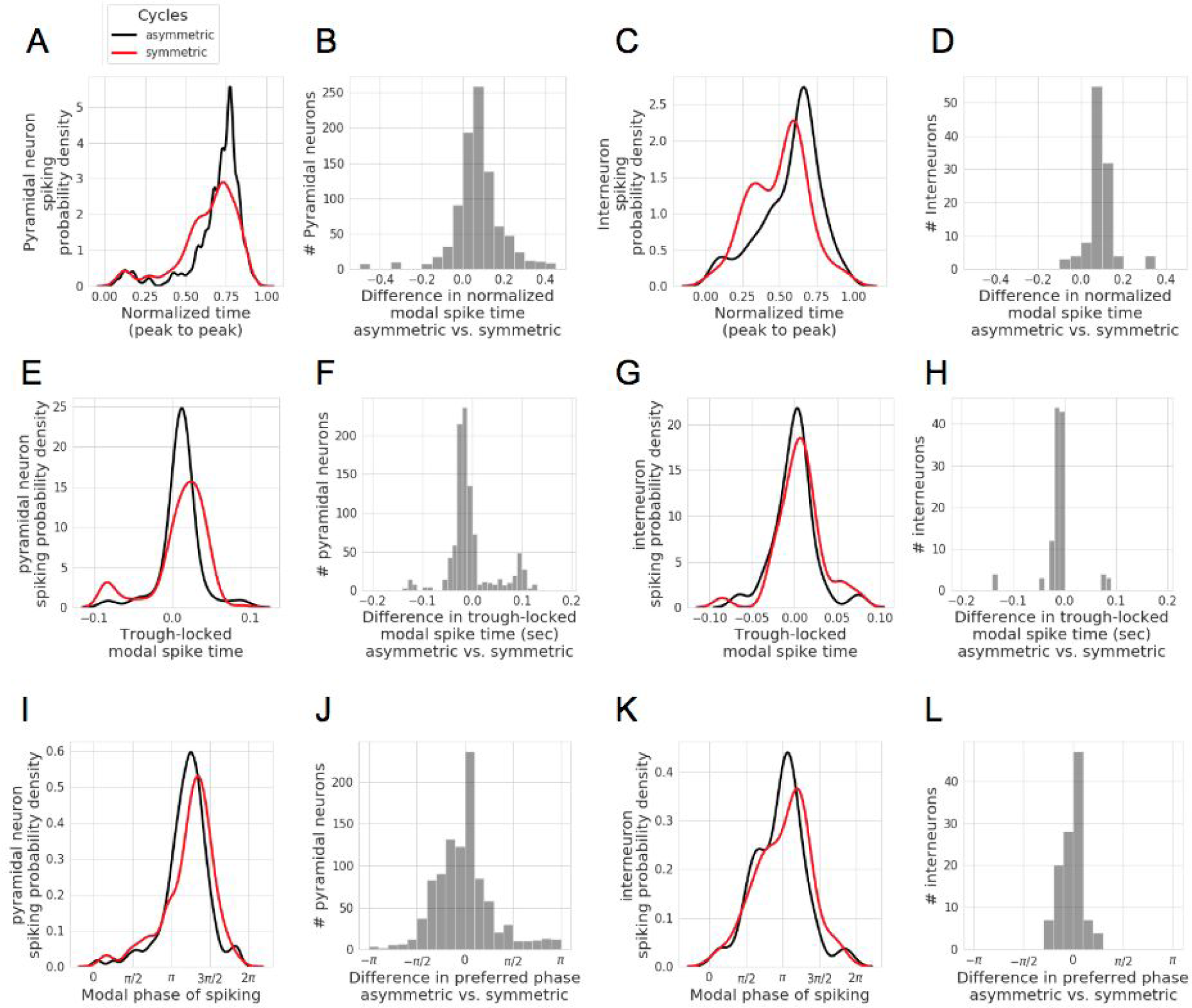
Analysis of spike timing changes with rise-decay symmetry. (A) Distributions of modal spike times across neurons. Time for each cycle was normalized such that 0 corresponded to the former peak, and 1 corresponded to the latter peak. Normalized spike times were collected during each cycle, and each cycle was defined as “asymmetric” or “symmetric” using a threshold on the rise-decay symmetry, and the modal spike time was computed (see Methods). The distribution shows the modal spike times across all putative pyramidal neurons during asymmetric cycles (black) and symmetric cycles (red). Note that the black distribution is shifted to the right of the red one, representing that neurons tended to fire at later normalized times during asymmetric cycles. (B) Distribution of the differences in modal normalized spike time between asymmetric and symmetric cycles for each neuron. Note that the distribution is shifted to the right of zero, showing that normalized spike times were systematically later during asymmetric cycles compared to symmetric cycles. (C-D) Same as A-B but for putative interneurons instead of pyramidal neurons. (E-H) Same as A-D but spike time was not normalized in each cycle, but rather referenced to the trough of the cycle. Note in (E) and (G) that the black (asymmetric cycles) distributions are shifted earlier in time, representing that neurons fire earlier relative to the trough during asymmetric cycles. This is also shown by the distributions in (F) and (H) that are shifted to the left of zero. (I-L) Same as A-D but spike phase was analyzed instead of spike time. Note in (I) and (K) that the black (asymmetric cycles) distributions are shifted earlier in the cycle, representing that neurons fire earlier in the cycle during asymmetric cycles. This is also shown by the distributions in (J) and (K) that are shifted to the left of zero.

However, we wondered if these timing trends could be more simply accounted for by imperfect extrema localization. We tested this in a simulation by generating a stationary oscillator with added brown noise (Figure S3A-C) so that the cycle-by-cycle characterization would not be able to exactly detect the “true” peaks and troughs of the underlying oscillatory generator, as is the case with experimental recordings. Spikes were generated randomly using a fixed spike-field coupling (phase-to-firing rate) mapping. Because of this noise, measured locations of extrema and cycle symmetries did not always match the ground truth. By analyzing spike timing in the same manner as described in the previous paragraph, we observed the same qualitative trends between spike timing and cycle asymmetry (Figure S3D-F), despite no differences in the spike-field coupling generator between asymmetric and symmetric cycles. Therefore, based on our results in Figure S2, we could not conclude that the symmetry of cycles indexes a change in neuronal spike timing, and we are not aware of a way to accurately assess the extent of the contribution of extrema localization noise. However, we have included a description of the analysis here to heed a warning to those who may explore similar analysis between waveform symmetry and spike timing.

**Figure S3.**
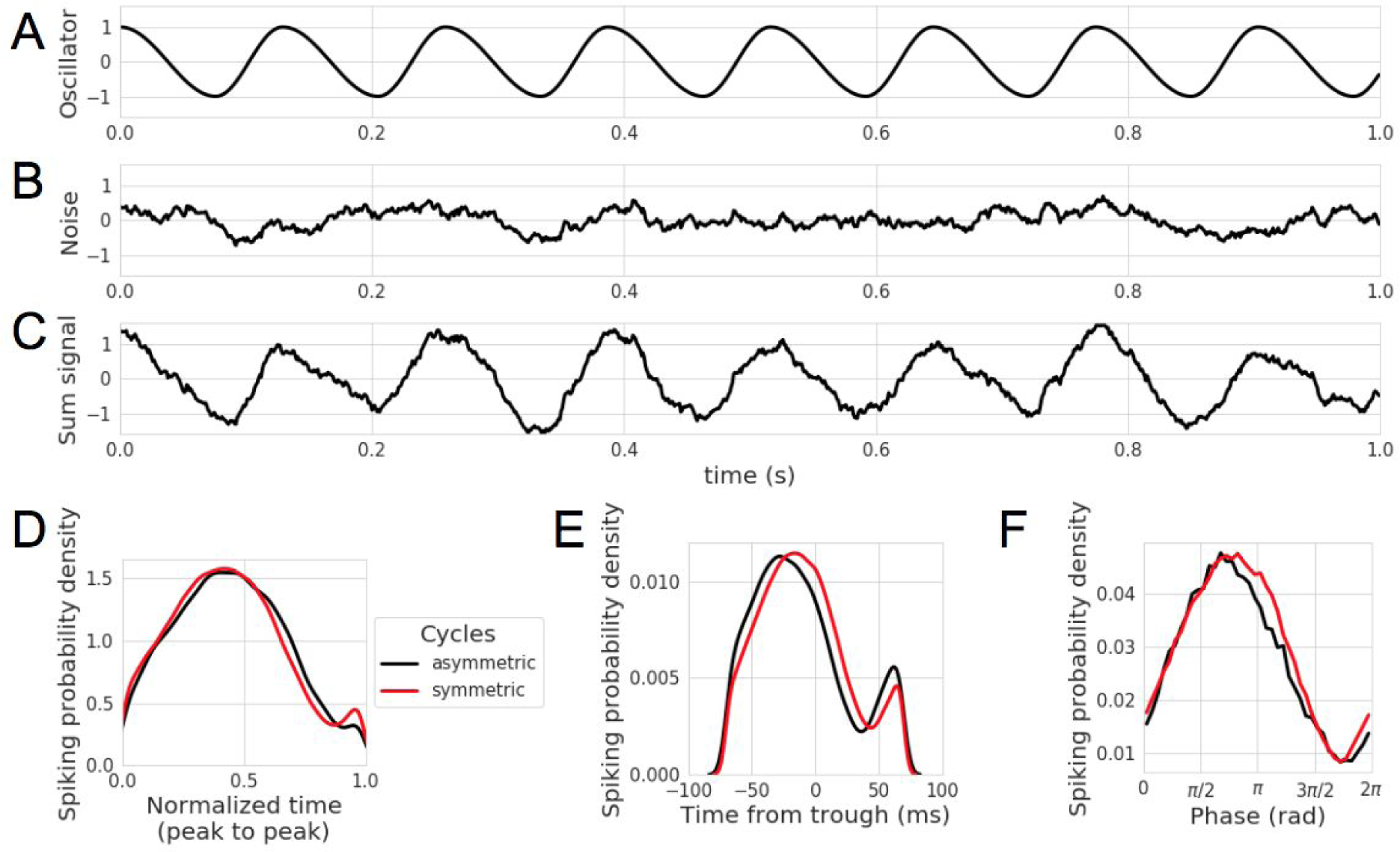
Spike timing and rise-decay symmetry correlations are an artifact of noise in extrema localization. (A-C) Simulating a noisy oscillation. (A) A stationary oscillation was simulated with a constant period and rise-decay symmetry which were the mean cycle features for an example hippocampal theta recording. (B) A brown noise process was generated and highpass filtered at 2Hz. (C) The components from (A) and (B) were summed together to generate a simulated signal containing an oscillation and noise. Note that the “ground truth” oscillator has a constant period and symmetry, but the cycle-by-cycle properties measured in the composite signal will have some variance due to the noise. (D-F) Spurious relationships between cycle symmetry and spike timing of a neuron that was simulated to spike with a stationary spike-field coupling relationship with respect to the ground-truth oscillation (A). These spurious relationships are due to uncertainties in the peak and trough localization caused by the noise process (B). Note in (D) that the asymmetric cycles have later normalized spike times compared to symmetric cycles (c.f. Figure S2 A,C). Note in (E) that the asymmetric cycles have earlier trough-relative spike times compared to symmetric cycles (c.f. Figure S2 E,G). Note in (F) that the asymmetric cycles have earlier spike phases compared to symmetric cycles (c.f. to Figure S2 I,K).

## Acknowledgements

We thank Thomas Donoghue, Richard Gao, Tammy Tran, and Roemer van der Meij for invaluable discussion. S.R.C. is supported by the National Science Foundation Graduate Research Fellowship Program and the University of California, San Diego Chancellor’s Research Excellence Scholarship. B.V. is supported by a Sloan Research Fellowship, the Whitehall Foundation (2017-12-73), and the National Science Foundation (1736028).

## Author contributions

S.R.C. performed the analyses. S.R.C. and B.V. designed the research and wrote the paper.

## Competing interests

The authors declare no competing interests.

